# High-Resolution Cryo-EM Structure Determination of α-Synuclein – A Prototypical Amyloid Fibril

**DOI:** 10.1101/2024.09.18.613698

**Authors:** Juan C. Sanchez, Josh Pierson, Collin G. Borcik, Chad M. Rienstra, Elizabeth R. Wright

**Author notes:** Elizabeth R. Wright **Email:**.

## Abstract

The physiological role of α-synuclein (α-syn), an intrinsically disordered presynaptic neuronal protein, is believed to impact the release of neurotransmitters through interactions with the SNARE complex. However, under certain cellular conditions that are not well understood, α-syn will self-assemble into β-sheet rich fibrils that accumulate and form insoluble neuronal inclusions. Studies of patient derived brain tissues have concluded that these inclusions are associated with Parkinson’s disease, the second most common neurodegenerative disorder, and other synuclein related diseases called synucleinopathies. In addition, repetitions of and specific mutations to the SNCA gene, the gene that encodes α-syn, results in an increased disposition for synucleinopathies. The latest advances in cryo-EM structure determination and real-space helical reconstruction methods have resulted in over 60 *in vitro* structures of α-syn fibrils solved to date, with a handful of these reaching a resolution below 2.5 Å. Here, we provide a protocol for α-syn protein expression, purification, and fibrilization. We detail how sample quality is assessed by negative stain transmission electron microscopy (NS-TEM) analysis and followed by sample vitrification using the Vitrobot Mark IV vitrification robot. We provide a detailed step by step protocol for high resolution cryo-EM structure determination of α-syn fibrils using RELION and a series of specialized helical reconstruction tools that can be run within RELION. Finally, we detail how ChimeraX, Coot, and Phenix are used to build and refine a molecular model into the high resolution cryo-EM map. This workflow resulted in a 2.04 Å structure of α-syn fibrils with excellent resolution of residues 36 to 97 and an additional island of density for residues 15 to 22 that had not been previously reported. This workflow should serve as a starting point for individuals new to the neurodegeneration and structural biology fields. Together, this procedure lays the foundation for advanced structural studies of α-synuclein and other amyloid fibrils.

**Key Features:**

- *In vitro* fibril amplification method yielding twisting fibrils that span several micrometers in length and are suitable for cryo-EM structure determination.
- High-throughput cryo-EM data collection of neurodegenerative fibrils, such as alpha-synuclein.
- Use of RELION implementations of helical reconstruction algorithms to generate high-resolution 3D structures of a-synuclein fibrils.
- Brief demonstration of the use of ChimeraX, Coot, and Phenix for molecular model building and refinement.s

**Graphical overview of α-synuclein fibrilization and cryo-EM structure determination:** α-synuclein protein expression and purification is followed by a fibrilization protocol yielding twisting filaments that span several micrometers in length and are validated by negative stain transmission electron microscopy (NS-TEM). The sample is then vitrified, followed by cryo-EM data collection. Real-space helical reconstruction is performed in RELION to generate an electron potential map that is used for model building.

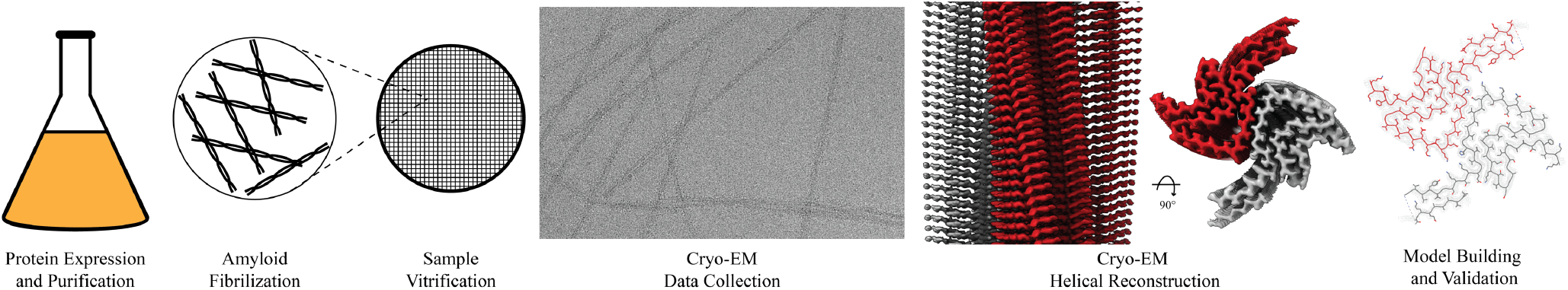

## Introduction and Background

Amyloid formation within neurons has been well documented to cause neurodegeneration in patients leading to a variety of diseases including Alzheimer’s (AH), Parkinson’s disease (PD), Lewy Body disease (LB), and multiple system atrophy (MSA) [1-3]. The formation of amyloids is due to protein aggregation resulting in helical, filamentous assemblies with cross β-sheet quaternary structure (Figure 1) [4]. Amyloid filaments interact with different cellular components such as membranes, cytoskeletal factors, and other filaments to form inclusion bodies that disrupt cellular processes and ultimately lead to cell death [2]. These inclusion bodies are prominent in postmortem brains of patients who have suffered from these neurodegenerative diseases, and early investigation of inclusion bodies revealed the presence of filamentous a-synuclein (α-syn) [1,2]. α-syn is a small (14.4 kDa) intrinsically disordered protein whose physiological role remains elusive. α-syn has the capability to bind to the SNARE complex and associate with vesicles at the neuronal axon terminus providing evidence that it may have an impact on neurotransmitter release, vesicle docking and vesicle trafficking [5-8]. However, upon misfolding, α-syn first forms oligomeric aggregates that eventually undergo fibrilization, these fibrils display the highly ordered cross β-sheets classically found in amyloids [9,10]. These, in turn, form the extended filaments that cause neuropathological changes in the brain and are specifically responsible for PD, LB, and MSA. Diseases caused by α-synuclein in this manner are called synucleinopathies [11].

**Figure 1.**
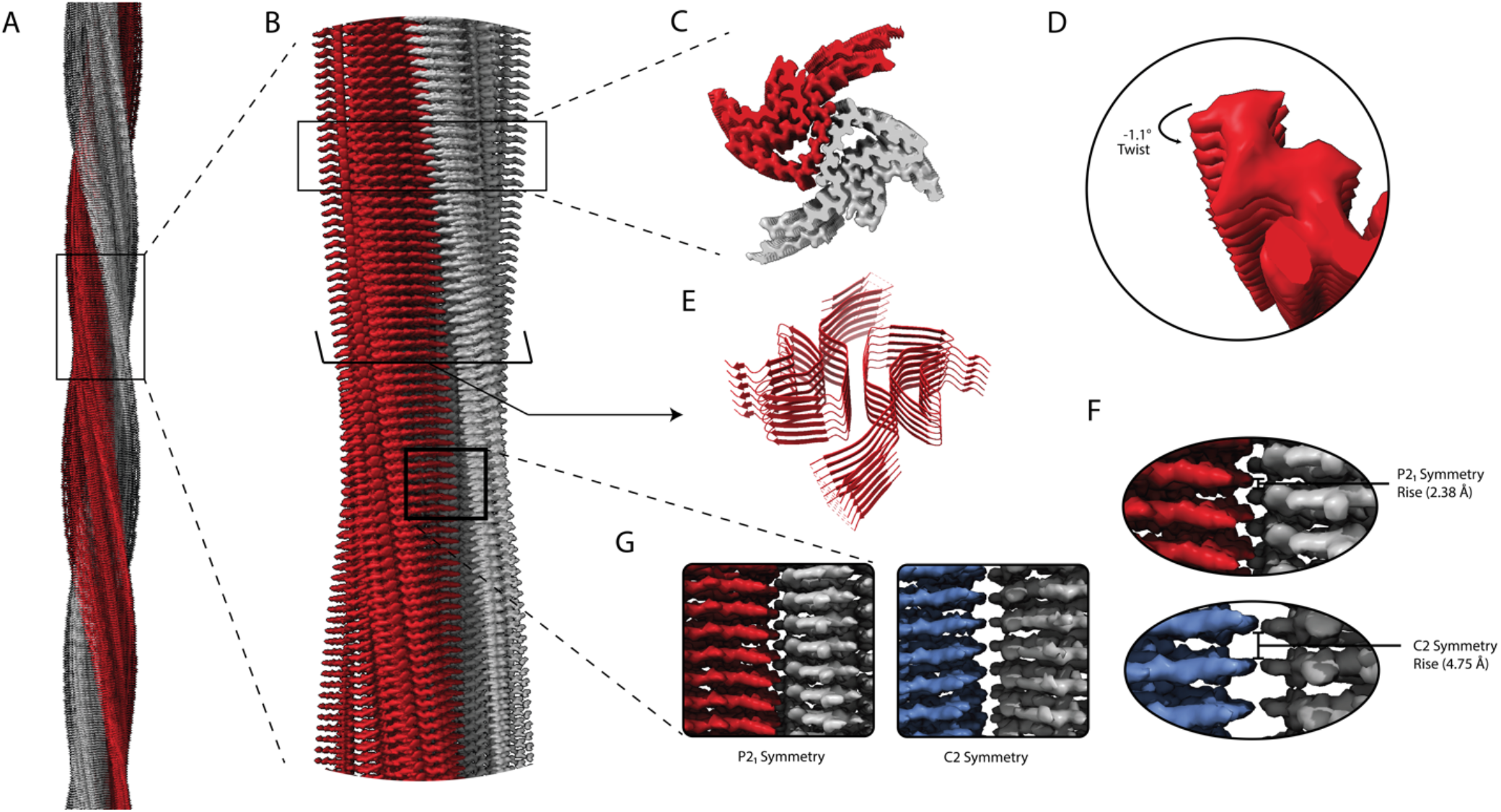
Structural features of α-syn fibrils from cryo-EM structures. A. Cryo-EM structure of full-length α-syn fibril depicting two protofilaments (one in red; one in grey). B. Magnified view of α-syn fibril portraying stacked rungs and filament twist. C. Cross-section of α-syn fibril electron potential map displaying two α-syn monomers that make up each protofilament approximately 180 degrees from each other. D. Electron potential map of individual β-sheet stacks twisting. E. Model depicting secondary structure of stacking β-sheets. F. Example of rise measurement for P2_1_ symmetry (Red) and C2 symmetry (Blue). G. Possible packing symmetry between protofilaments for P2_1_ symmetry (out of register) (Red) and C2 symmetry (in register) (Blue).

The high-resolution structure presented here of filamentous wild-type α-syn is of a helical filament composed of 2 protofilaments and each turn (or rung) of the filament is comprised of 2 copies (one per protofilament) of α-syn facing nearly 180 degrees from each other (Figure 1). Between the monomers that make up each protofilament there is a hydrophobic interface composed of residues 50-57, similar to previously solved structures of filamentous α-syn [12-14]. This interface is stabilized by salt-bridges and pseudo screw symmetry, as previously reported [12,13]. For α-syn, there are 7 different missense familial mutations commonly found in patients who have a higher disposition for synucleinopathies (A30P, E46K, H50Q, G51D, A53E, A53T, and A53V) [15-21]. Interestingly, 6 of these familial mutations lie within the core of the structure and may cause destabilization resulting in a variety of different fibril morphologies. The presence of polymorphism has been demonstrated particularly well through the analysis of *in vitro* α-syn fibrils. Fibril twist, crossover distance, packing arrangement, number of protofilaments, interface, tertiary structure, etc. can vary greatly under different micro- and macro-environments. Many different environmental factors such as pH, salt concentrations, temperature, quiescence, and post translational modifications have an impact on fibril morphology—this has led to documentation of more than 60 *in vitro* structural polymorphs of α-syn in the PDB [22,23]. These structural differences in the *in vitro* filaments can have direct effects on nucleation rates, seeding propensities, and even cytotoxicity [23]. Unfortunately, the ties between these structurally distinct *in vitro* polymorphs to those found in sarkosyl-insoluble brain-derived structures remains elusive. However, evidence suggests that different polymorphs may influence pathologies [24-26]. This is demonstrated by the difference in α-syn folds of the filaments extracted from patients diagnosed with MSA versus PD [27].

The formation of the filaments responsible for synucleinopathies are propagated in brain tissue by primary nucleation events in which α-syn monomer spontaneously undergoes structural changes resulting in nucleation. This nucleation site can then recruit additional α-syn monomers to bind, thus elongate the fibril [28,29]. However, there can also be secondary nucleation events in which preformed fibrils are introduced into the cellular environment as “seeds” [30]. These seeding events are significantly more potent at fibril formation and elongation. Remarkably, seeds from a particular polymorph have been shown to recruit wild-type α-syn, provide a structural template, and form filaments expressing the polymorph of the seed regardless of whether the endogenous protein recruited is pathogenic or not [31]. A consequence of this prion-like self-replication is that α-syn fibrils may move from cell-to-cell spreading cytotoxic polymorphs.

The introduction of polymorphism has a multifactorial effect on clinical treatments of neurodegenerative diseases. Our understanding of the implications associated with each polymorph on disease progression, pathology, and patient outcomes is very limited. In addition, the differences in folding, packing, twists, etc. of each polymorph introduces complexities in binding sites, affinities, and accessibility for a “one size fits all” drug for synucleinopathies; this is further complicated by evidence that not only are there disease specific morphisms, but evidence shows that each synucleinopathy can exhibit patient-to-patient heterogeneity [32]. Thus, to overcome these challenges, explore new therapeutic targets, understand specific polymorph effects on neuropathology, and develop therapies with patient-specific approaches, solving both patient-derived and *in vitro* amyloid polymorphs should be explored.

Here, we describe a helical reconstruction workflow that we use to solve the structure of *in vitro* assembled filamentous α-syn to a global resolution of 2.04 Å. We purify α-syn filaments from a reaction in which fibril seeding material is combined with monomeric α-syn. The fibril seeding material provides a template for fibril elongation via monomer addition over a 6-week incubation period at 37 °C with shaking at 200 rpm. The purified α-syn filaments are then imaged using negative stain transmission electron microscopy (NS-TEM) to evaluate sample integrity and fibril concentration on the grid. The sample is then applied to grids and plunge frozen, and the vitrified grids are used for cryo-EM data collection. We provide a detailed protocol utilizing RELION to reconstruct a high-resolution cryo-EM electron potential map that is then used for building an atomic model of the fibril (Figure 1B, 1C, 1E). The steps presented here may be applied to studies of various amyloid fibrils and accelerate cryo-EM structure determination in the fields neurodegenerative research and medicine.

## Materials and Reagents

### Biological materials

1. Plasmid with wild-type α-syn construct in *E*. *coli* BL21(DE3)/pET28A-AS [33].

### Reagents

1. LB broth (Invitrogen, catalog number: 12780029)
2. Bacto Agar (Dot Scientific Inc., catalog number: DSA20030-1000)
3. Magnesium sulfate, MgSO_4_ (Fisher Scientific, catalog number: 01-337-186)
4. Calcium chloride, CaCl_2_ (Fisher Scientific, catalog number: BP510-500)
5. Sodium phosphate, NaH_2_PO_4_ (Fisher Scientific, catalog number: 01-337-702)
6. Potassium phosphate, KH_2_PO_4_ (Fisher Scientific, catalog number: 01-337-803)
7. Sodium chloride, NaCl (Fisher Scientific, catalog number: S271-500)
8. IPTG (Fisher Scientific, catalog number: BP1755-10)
9. Tris-HCl (Fisher Scientific, catalog number: PRH5125)
10. EDTA (Fisher Scientific, catalog number: AAA1516130)
11. Kanamycin monosulfate (Thermo Scientific, catalog number: J61272.14)
12. SDS-PAGE gels (Bio-Rad, catalog number: 4561096)
13. SDS-PAGE Loading Dye (Bio-Rad, catalog number: 1610737)
14. Coomassie Brilliant Blue (TCI, catalog number: 6104-59-2)
15. BME vitamins (Sigma-Aldrich, catalog number: B6891-100mL)
16. Sodium azide (Sigma-Aldrich, catalog number: 19-993-1)
17. Studier trace metal mix (Sigma-Aldrich, catalog number: 41106212)
18. Ammonium Sulfate (Fisher Scientific, catalog number: A702-500)
19. Deuterium oxide, ^2^H_2_O (Cambridge Isotopes Laboratories, catalog number: DLM-4-1L)
20. BioExpress Bacterial Cell Media 10X concentrate (U-^13^C, 98%; U-^15^N, 98%; U-D 98%) (Cambridge Isotopes Laboratories, catalog number: CGM-1000-CDN)
21. ^15^N-NH_4_CI (Cambridge Isotopes Laboratories, catalog number: 39466-62-10)
22. ^2^H-^13^C-glucose (Cambridge Isotopes Laboratories, catalog number: CDLM-3813-5)
23. Sodium deuteroxide, NaO^2^H (Cambridge Isotopes Laboratories, catalog number: DLM-45-100)
24. 2% Uranyl Acetate (UA) (EMS, catalog number: 22400-2)

### Solutions

1. Kanamycin Stock Solution (1000x, 40 mg/ml) (recipe below)
2. Kanamycin Stock Solution (1000x, 90 mg/ml) (recipe below)
3. Conditioning Plate (recipe below)
4. Pre-Growth Media (recipe below)
5. Wash Buffer (recipe below)
6. Growth Media (recipe below)
7. IPTG Stock Solution
8. Buffer A (recipe below)
9. Buffer B (recipe below)
10. TEN Buffer (recipe below)
11. Saturated Ammonium Sulfate Solution (recipe below)
12. Fibrilization Buffer (recipe below)
13. 1% Uranyl Acetate (recipe below)

### Recipes

1. Kanamycin Stock Solution (1000x, 40 mg/ml)

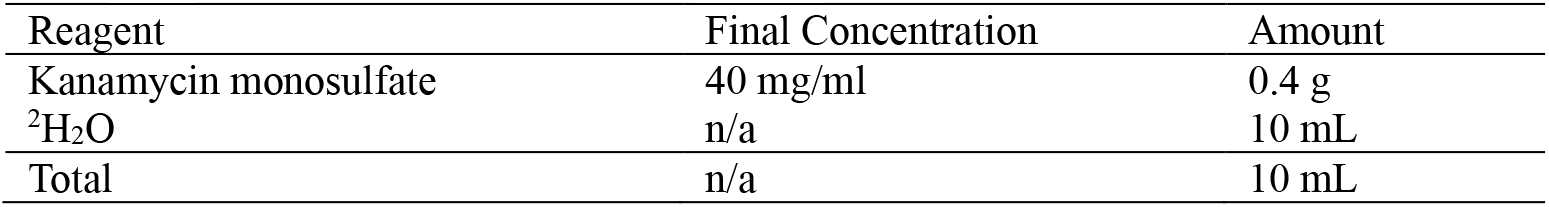
  1. Completely dissolve kanamycin monosulfate in ^2^H_2_O
  2. Sterilize solution using a 0.22 μm syringe filter (GenClone) and 10 mL syringe (BD)
  3. Aliquot 1000 uL stocks and store at -20°C until use.
2. Kanamycin Stock Solution (1000x, 90 mg/ml)

  1. Completely dissolve kanamycin monosulfate in ^2^H_2_O
  2. Sterilize solution using a 0.22 μm syringe filter (GenClone) and 10 mL syringe (BD)
  3. Aliquot 1000 uL stocks and store at -20°C until use.
3. Conditioning Plate

| Reagent | Final Concentration | Amount |
| --- | --- | --- |
| Kanamycin monosulfate | 90 mg/ml | 0.9 g |
| <sup>2</sup>H<sub>2</sub>O | n/a | 10 mL |
| Total | n/a | 10 mL |

  1. Combine reagents in a flask and autoclave at 121°C, 15 PSI for at least 20 minutes.
  2. Allow the media to cool to ∼55°C, then add 1000 uL of the kanamycin stock solution (1000x, 40 mg/ml)
  3. Pour ∼25 mL of media per Petri plate (100 mm), repeat for remaining 1 L.
4. Pre-Growth Media

| Reagent | Final Concentration | Amount |
| --- | --- | --- |
| <sup>2</sup>H<sub>2</sub>O | 70% | 700 mL |
| LB Broth | 2% | 20 g |
| Bacto Agar | 1.5% | 15 g |
| H<sub>2</sub>O | n/a | Fill to 1000 mL |
| Total | n/a | 1000 mL |

  1. Combine reagents in a flask and autoclave at 121°C, 15 PSI for at least 20 minutes.
  2. Allow the media to cool to ∼55°C, then add 50 uL of the kanamycin stock solution (1000x, 40 mg/ml)
5. Wash Buffer

| Reagent | Final Concentration | Amount |
| --- | --- | --- |
| <sup>2</sup>H<sub>2</sub>O | 70% | 35 mL |
| LB Broth | 2% | 1 g |
| H<sub>2</sub>O | n/a | Fill to 50 mL |
| Total | n/a | 50 mL |

  1. Combine reagents and pH to 7.6 with NaO^2^H (Cambridge Isotope Laboratories)
  2. Sterile filter solution using a 50 mL filtration system (Steriflip)
6. Growth Media

| Reagent | Final Concentration | Amount |
| --- | --- | --- |
| NaH<sub>2</sub>PO<sub>4</sub> | 50 mM | 0.34 g |
| KH<sub>2</sub>PO<sub>4</sub> | 25 mM | 0.17 g |
| NaCl | 10 mM | 0.03 g |
| <sup>2</sup>H<sub>2</sub>O | n/a | Fill to 50 mL |
| Total | n/a | 50 mL |

  1. Combine reagents and add 1000 uL of the kanamycin stock solution (1000x, 90 mg/ml), and pH to 7.6 with NaO^2^H (Cambridge Isotope Laboratories)
  2. Sterile filter solution using a 1000 mL filtration system (Fisher Scientific)
7. IPTG Stock Solution

| Reagent | Final Concentration | Amount |
| --- | --- | --- |
| NaH<sub>2</sub>PO<sub>4</sub> | 50 mM | 6.9 g |
| KH<sub>2</sub>PO<sub>4</sub> | 25 mM | 3.4 g |
| NaCl | 10 mM | 0.58 g |
| MgSO<sub>4</sub> | 5 mM | 1.23 g |
| CaCl<sub>2</sub> | 0.2 mM | 0.03 g |
| Bacterial Cell Media 10x | 0.1x | 10 mL |
| BME vitamins 100x | 0.25x | 2.5 mL |
| Studier trace metals 1000x | 0.25x | 0.25 mL |
| <sup>15</sup>N-NH<sub>4</sub>Cl | 1 g/L | 1 g |
| <sup>2</sup>H-<sup>13</sup>C-glucose | 8 g/L | 8 g |
| BME vitamins | 0.25x | 2.5 mL |
| <sup>2</sup>H<sub>2</sub>O | n/a | Fill to 50 mL |
| Total | n/a | 1000 mL |

  1. Completely dissolve IPTG in ^2^H_2_O
  2. Sterilize solution using a 0.22 μm syringe filter (GenClone) and 10 mL syringe (BD)
  3. Aliquot 1000 uL stocks and store at -20°C until use.
8. Buffer A

| Reagent | Final Concentration | Amount |
| --- | --- | --- |
| IPTG | 0.5 M | 1.2 g |
| <sup>2</sup>H<sub>2</sub>O | n/a | 10 mL |
| Total | n/a | 10 mL |

  1. Dissolve Tris-HCL and NaCl in water while stirring.
  2. pH buffer to 7.4 at 37°C using 1M NaOH.
9. Buffer B

| Reagent | Final Concentration | Amount |
| --- | --- | --- |
| Tris-HCL | 30 mM | 4.73 g |
| NaCl | 30 mM | 1.75 g |
| H<sub>2</sub>O | n/a | 1 L |
| Total | n/a | 1L |

  1. Dissolve Tris-HCL and NaCl in water while stirring.
  2. pH buffer to 7.4 at 37°C using 1M NaOH.
10. TEN Buffer

| Reagent | Final Concentration | Amount |
| --- | --- | --- |
| Tris-HCL | 30 mM | 2.36 g |
| NaCl | 1 M | 29.22 g |
| H<sub>2</sub>O | n/a | 500 mL |
| Total | n/a | 500 mL |

  1. Dissolve Tris-HCL and NaCl in water while stirring.
  2. pH buffer to 7.4 at 37°C using 1M NaOH.
11. Saturated Ammonium Sulfate Solution

| Reagent | Final Concentration | Amount |
| --- | --- | --- |
| Tris-HCL | 30 mM | 4.73 g |
| NaCl | 30 mM | 1.75 g |
| EDTA | 0.1 mM | 29.22 mg |
| H<sub>2</sub>O | n/a | 1 L |
| Total | n/a | 1L |

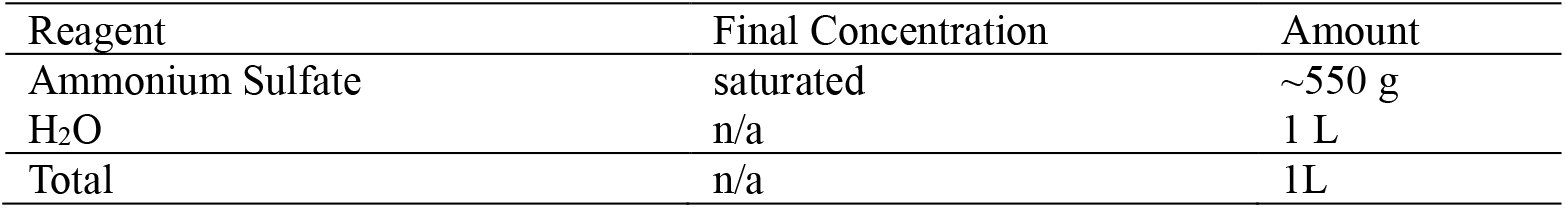
  1. Add ammonium sulfate into water while stirring.
  2. Heat gently until all ammonium sulfate is dissolved.
  3. Cool to room temperature. Crystals should form to indicate the solution is saturated.
12. Fibrilization Buffer

  1. Add NaH_2_PO_4_, EDTA, 0.02% sodium azide solution into ^2^H_2_O and H_2_O.
  2. pH to 7.4 at 37°C using 1M NaO^2^H.
13. 1% Uranyl Acetate (UA)

| Reagent | Final Concentration | Amount |
| --- | --- | --- |
| NaH<sub>2</sub>PO<sub>4</sub> | 50 mM | 1.2 g |
| EDTA | 0.1 mM | 5.85 mg |
| Sodium Azide | 0.02% | 40 ug |
| H<sub>2</sub>O | 20% | 40 mL |
| <sup>2</sup>H<sub>2</sub>O | 80% | 160 mL |
| Total | n/a | 200 mL |

  1. Mix 1-part sterile water with 1-part 2% UA stain (EMS) and filter through a Spin-X centrifuge tube with a 0.22 μm filter (Costar).

13. 1% Uranyl Acetate (UA)
| Reagent | Final Concentration | Amount |
| --- | --- | --- |
| 2 % Uranyl Acetate | 1% (v/v) | 250 µL |
| H<sub>2</sub>O | n/a | 250 µL |
| Total | n/a | 500 µL |

### Laboratory Supplies

1. 10 ml Syringe (BD, catalog number: 309604)
2. 0.22 um filter (GenClone, catalog number: 25-240)
3. 100 mm × 15 mm Petri dishes (Fisher Scientific, catalog number: S33580A)
4. 1000 mL filtration system (Fisher Scientific, catalog number: FB12566506)
5. 50 mL conical tubes (VWR, 525-1074)
6. 0.45 μm syringe filter (GenClone, catalog number: 25-246)
7. 1.7 mL centrifuge tubes (Denville, catalog number: C2170)
8. Parafilm (Bemis, catalog number: PM996)
9. 0.22 μm Spin-X centrifuge tube filter (Costar, catalog number: 8160)
10. 200 mesh carbon film, copper grids (EMS, catalog number: CF200-CU)
11. Whatman #1 filter paper (Whatman, catalog number: 1001-090)
12. Quantifoil R2/1 200 mesh, copper grids (Quantifoil Micro Tools GmbH, catalog number: Q210CR1)
13. Standard Vitrobot Filter Paper, Ø55/20mm, Grade 595 (Ted Pella, catalog number: 47000-100)

### Equipment

1. HiTrap Q HP anion exchange column (Cytiva, catalog number: 17115401)
2. Stirred cell concentrator (Amicon, catalog number: UFSC05001)
3. HiPrep 16/60 Sephacryl S100-HR gel filtration column (Cytiva, catalog number: 17119501)
4. 5424 R Microcentrifuge (Eppendorf, catalog number: 05-400-005)
5. Grid holder block (Pelco, catalog number: 16820-25)
6. Plasma Cleaner (Harrick Plasma Inc., catalog number: PDC-32G)
7. Static dissipator (Mettler Toledo, catalog number: UX-11337-99)
8. Style N5 reverse pressure tweezers (Dumont, catalog number: 0202-N5-PS-1)
9. Talos L120C 120 kV transmission electron microscope (TEM) (Thermo Fisher Scientific), or equivalent
10. Cryo grid box (Sub-Angstrom, catalog number: SB)
11. Plasma Cleaner (Harrick Plasma Inc., catalog number: PDC-32G)
12. Vitrobot Mark IV vitrification robot (Thermo Fisher Scientific)
13. Titan Krios G3i 300 kV transmission electron microscope (TEM) (Thermo Fisher Scientific)
14. K3-GIF direct electron detector with energy filter (Gatan Inc., AMETEK)
15. High-performance computing (HPC) cluster with an EPYC Milan 7713P 64-core 2.0GHz CPU (AMD), 512 GB RAM, 4x RTX A5000 24GB GDDR6 GPU (NVIDIA), 2x 960GB Enterprise SSD, mirrored OS, 2x 7.68TB nVME SSD as 15TB scratch space, dual-port 25GbE Ethernet.

### Software

1. SBGrid (https://sbgrid.org/)
2. IMOD (https://bio3d.colorado.edu/imod/)
3. RELION (https://relion.readthedocs.io/en/release-4.0/)
4. MotionCor2 (https://emcore.ucsf.edu/ucsf-software)
5. Gctf (https://sbgrid.org/software/titles/gctf)
6. Topaz-filament (https://github.com/3dem/topaz)
7. UCSF ChimeraX (https://www.cgl.ucsf.edu/chimerax/)
8. Coot (https://www2.mrc-lmb.cam.ac.uk/personal/pemsley/coot/)
9. PHENIX (https://phenix-online.org/)

## Procedure and Results

### A. α-synuclein sample preparation

Expression and purification of α-syn protein is performed as reported previously [33]. The protein preparations and fibrilization protocol presented here were developed for joint cryo-EM and NMR studies. Preparations include the use of isotopically labeled reagents that are critical for NMR experiments but are not necessary for cryo-EM. Thus, the α-syn sample preparation protocol may be adapted for cryo-EM only studies by substituting isotopically labeled reagents with a standard equivalent reagent.

#### α-synuclein protein expression

1. Expression of wild-type α-syn is performed in *E. coli* BL21(DE3)/pET28a-AS.
2. Plate transformed cells onto conditioning plate, overnight at 37 °C.
3. Inoculate a 50 mL pre-growth flask with a single colony from the overnight conditioning plate and incubate overnight at 220 rpm at 37 °C until OD_600_ = ∼3.
4. Transfer cells into a 50 mL conical tube (VWR) using aseptic techniques. Centrifuge tubes at 5000 rpm for 5 minutes at 4 °C. Decant supernatant and wash with ∼20 mL of cold wash buffer.
5. Resuspended cells with the growth media in 4x 1L baffled flasks, 250 mL each. At an OD_600_ of ∼1-1.2, induce α-syn over-expression with the addition of 1 mL of an IPTG stock solution. Incubate at 25 °C with shaking at 200 rpm.
6. After overnight growth, collect cells and combine for harvesting (∼15 hours post-induction). Centrifuge at 5000 rpm for 10 minutes at 4 °C. Decant the supernatant and wash the cell pellet with the wash buffer to remove residual growth media components.
7. Cell pellets may then be frozen and stored at -80 °C until use.

#### α-synuclein protein purification

1. Cells may be lysed via heat denaturation, as α-syn is thermostable and will be unaffected. Place 50 mL conical tubes (VWR) containing cell paste in boiling water (98°C) for 30 minutes. Cool cell lysate on ice. Clear the cell lysate by centrifugation at 5000 rpm for 10 minutes at 4 °C.
2. α-syn should then be precipitated via addition of a saturated ammonium sulfate solution on ice. Collect α-syn precipitate via centrifugation at 16900 rpm for 45 minutes at 4 °C and decanting the supernatant.
3. Equilibrate the HiTrap Q HP anion exchange column (Cytiva) with Buffer A.
4. Resolubilize the α-syn precipitate with ∼5 ml Buffer A. Make sure to filter the resolubilized α-syn using a 0.45 μm syringe filter (GenClone). Inject the resolubilized α-syn to bind to the QFF anion exchange resin (GE Healthcare Life Sciences, Marlborough, MA). Elute using a linear gradient of 0.03–0.6 M NaCl by increasing the proportion of Buffer B flow through the column. Collect fraction as they come off the column. In our hands, fractions containing α-syn monomer usually elute at about 0.3 M NaCl.
5. After completion, Run SDS-PAGE to check to determine which fractions (gel bands) contain α-syn. Take 20 μL samples from each fraction tube from 20% Buffer B to 40% Buffer B. Add 20 μL 2x SDS loading dye (Bio-Rad) to each sample tube and heat at 90°C for 5 minutes. Run all samples on an SDS-PAGE gel (Bio-Rad). Use Coomassie Brilliant Blue stain (Sigma-Aldrich) to stain the gel. Examine the stained gel for α-syn over-expression bands. Note that α-syn tends to run at an apparent size of 18 kDa. Pool these fractions.
6. Concentrate the α-syn monomer solution using a stirred cell concentrator (Amicon) using a 3.5 kDa molecular weight cut off filter to a final concentration of ∼15 mg/mL. Prewet the concentrator with Buffer A before adding α-syn solution to prevent loss of sample to the filter.
7. Equilibrate the 16/60 Sephacryl S-200 HR gel filtration column (GE Healthcare Life Sciences) with TEN Buffer, 5x column volume.
8. Inject 1 mL of the concentrated α-syn pool into the loop path of the 16/60 Sephacryl S-100 HR gel filtration column (Cytiva) and run the protocol at 0.5 mL/min until the fraction with an apparent mass of 15 kDa, at ∼97 minutes.
9. Pool fractions, concentrate to ∼15 mg/ml α-syn using a clean stirred cell concentrator (Amicon) and a 3.5 kDa molecular weight cut off filter. Prewet the unit and filter with TEN buffer before adding α-syn solution to prevent loss of sample to the filter.
10. Purified α-syn may then be frozen and stored in -80 °C freezer until use.

#### α-synuclein fibrilization

1. Buffer exchange from the TEN buffer to fibrilization buffer. Add purified α-syn from above to a prewetted (with the fibrilization buffer) stirred cell concentrator (Amicon) and a 3.5 kDa molecular weight cut off filter. Dilute 10x with fibrilization buffer and concentrate down to the initial volume. Repeat 3 times to effectively remove TEN buffer and completely exchange to fibrilization buffer.
2. Purified α-syn protein in above buffer was concentrated to 15 mg/ml using 3.5 kDa cut off stir cell concentrators and 0.5 ml aliquoted into clean, sterile 1.7 ml Eppendorf tubes (Denville).
3. Fibril formation may be seeded with ∼50 ng of previously made mature α-syn fibril (in this case: sample used to determine the PDB ID: 2N0A fibril structure).
4. Seal the tubes with parafilm (Bermis) for the duration of the incubation.
5. Incubate at 37 °C and shake at 250 rpm continuously for 3 weeks. The viscosity of the fibril solution will greatly increase over time.
6. At the end of 3 weeks, add 100 μL of fibrilization buffer and continue the incubation for 3 weeks under the same conditions.
7. After a total of 6 weeks the fibrils at a protein concentration of ∼13 mg/ml are ready for TEM analysis.

### B. Negative stain

Fibrilization can be characterized by thioflavin-T (ThT) assays, which leverage the fluorescence signal observed when thioflavin-T binds to fibrils, a property not observed in the presence of purified protein monomers [34]. Although this method is powerful and can even detail fibrilization kinetics, there are limitations in the technique. Specifically, this assay can not specify whether fibrils are twisting, if the fibrils span several micrometers in length, or are small fragments 10s of nanometers in length. For high-resolution cryo-EM structure determination, fibrils should be both twisting and span several crossovers. Additionally, fibrils should be concentrated to a point where several fibrils span the micrograph but are not crowded or overlapping. This ensures there are enough individual particles for the reconstruction process. To determine if the fibrils possess these qualities, we perform negative stain transmission electron microscopy (NS-TEM) with the following procedure to test a range of sample concentrations. We found that a concentration of 6.5 mg/ml (i.e., 1:1 ratio of sample to buffer) was best for our sample on the grid.

1. Place the desired number of 200 mesh carbon film, copper EM grids (EMS) on a grid holder block (Pelco) and using a plasma cleaner PDC-32G (Harrick Plasma Inc.), or equivalent system, to glow discharge grids under a 100-micron vacuum for 30 seconds on low (Figure 2, step 1).
2. Cut a piece of parafilm (Bemis) to about 2”x4” and demagnetize with a static dissipater (Mettler Toledo) (Figure 2, step 2).
3. Retrieve one glow discharged EM grid using style N5 reverse pressure tweezers (Dumont), or similar tweezers (Figure 2, step 3).
4. Spot two 50 μL drops of sterile, Nanopure water and two 50 μL drops of 1% UA on to the piece of parafilm (Bemis), ensure the drops do not touch (Figure 2, step 4).
5. Apply 4 μL of the sample to the EM grid and allow the sample to incubate at room temperature for one minute (Figure 2, step 5).
6. Blot away the liquid by touching the edge of the EM grid to a piece of filter paper (Whatman) (Figure 2, step 6).
7. Wash the EM grid by touching the face of the EM grid to the 1^st^ drop of water, then blot away the liquid as in step 6. Repeat, but this time wash with the 2^nd^ drop of water (Figure 2, step 7).
8. Pre-stain the EM grid by touching the face of the EM grid to the 1^st^ drop of 1% UA, then blot away the stain as in step 6 (Figure 2, step 8).
9. Stain the grid by holding the face of the EM grid to the 2^nd^ drop of 1% UA for 15 seconds, then blot away the stain as in step 6 (Figure 2, step 9).
10. Allow the EM grid to dry for at least 5 minutes at room temperature before storing the grid in a grid box (Figure 2, step 10). Store the grid box in a desiccator or humidity-controlled room until imaging.
11. Repeat for additional sample dilutions to assess the sample conditions that may be best suited for cryo-EM analysis. We imaged the sample at a concentration of 13 mg/ml (undiluted), 6.5 mg/ml (2x dilution), and mg/ml (5x dilution) and found that a concentration of 6.5 mg/ml showed the best sample distribution on the grid (Figure 3). Note: Since fibrilization conditions greatly impact the length of the fibrils, and thus sample distribution on the grid, it is important to test each sample by NS-TEM before sample vitrification and cryo-EM data collection.
12. Image grids on a Talos L120C 120 kV TEM, or equivalent microscope, at a pixel size of 1.58 Å and a total electron dose of 25 e^-^/Å^2^.

**Figure 2.**
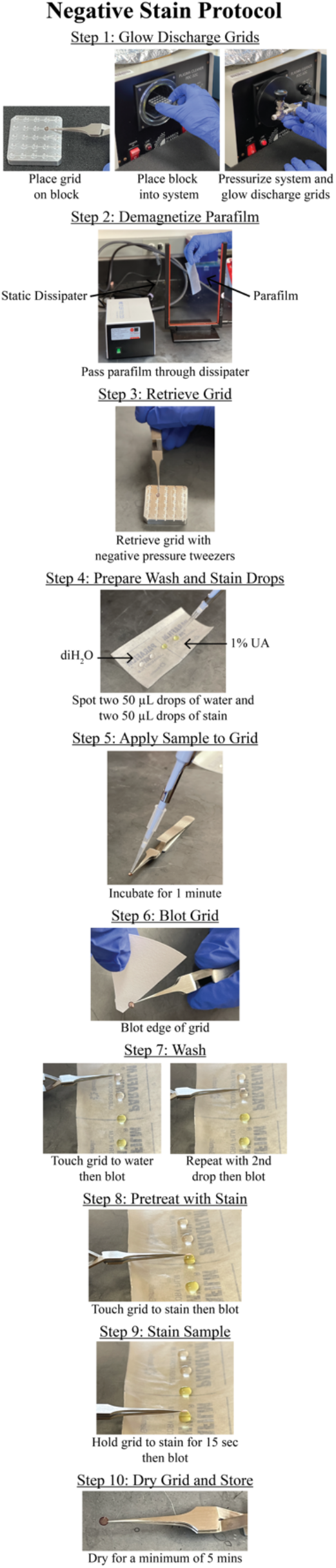
Negative stain protocol. Detailed steps for preparing negative stain grids of α-syn fibrils. The protocol yields lightly stained fibrils allowing for the visualization twisting fibrils comprised of two protofilaments (Figure 3). The procedure is repeated spanning a range of fibril concentrations that are imaged by transmission electron microscopy.

**Figure 3.**
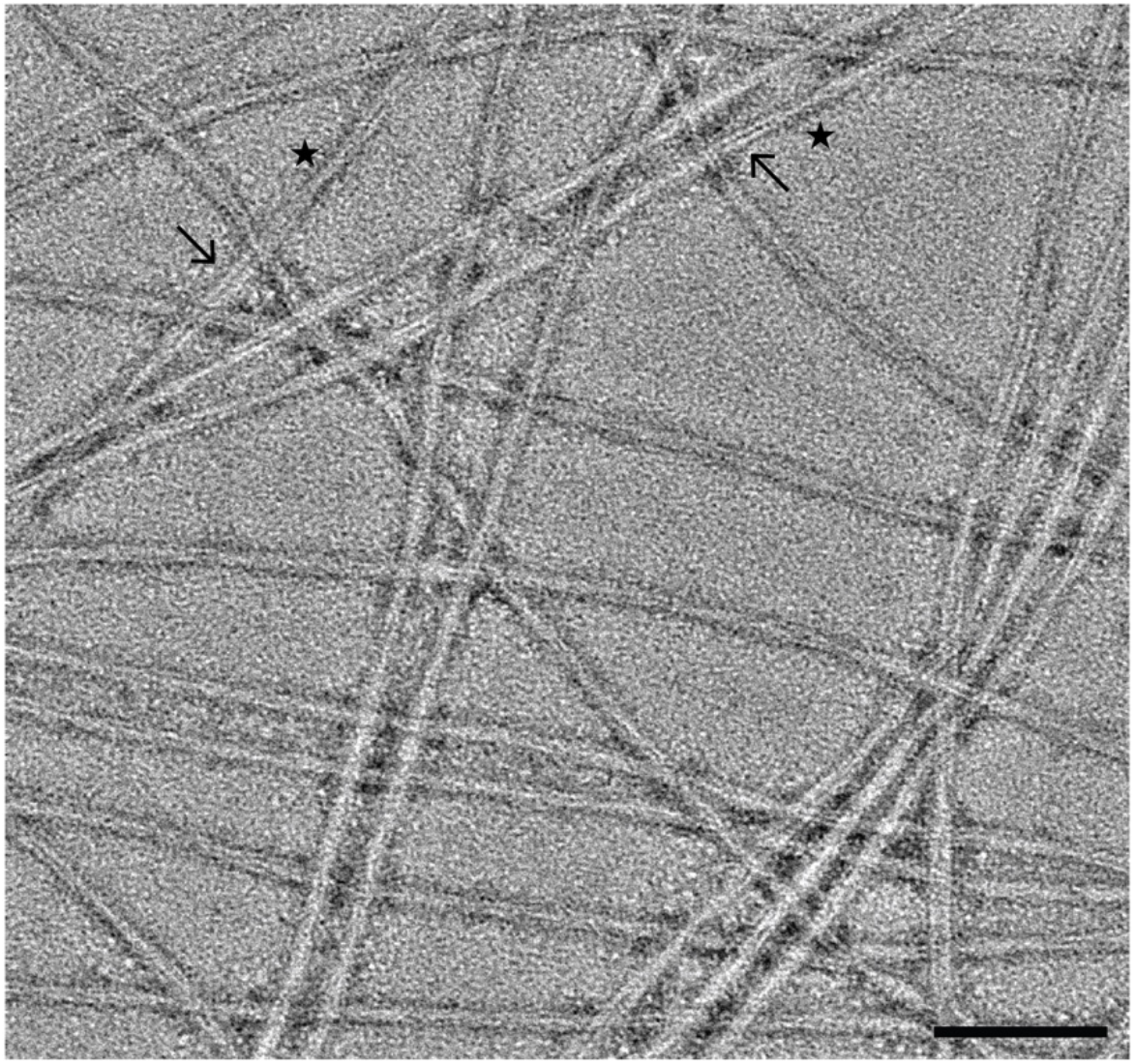
Negative stain TEM analysis of α-synuclein fibrils. Representative micrograph of fibrils lightly stained with 1% UA. The fibrils are comprised of two protofilaments (arrows) and appear to be twisting with distinct crossover points (stars). Scale bar, 100 nm.

### C. Sample vitrification

Basic sample vitrification for single particle analysis has become routine in the cryo-EM field. Here, we present a brief workflow of the vitrification process using the Vitrobot Mark IV (Thermo Fisher Scientific) with blotting conditions that yield grids suitable for cryo-EM data collection.

1. Using a plastic syringe (BD), add 60 mL of distilled water to the Vitrobot Mark IV water reservoir.
2. Turn on the Vitrobot Mark IV, set the chamber temperature to 20°C and the relative humidity to 95%.
3. Attach standard Vitrobot filter paper (Ted Pella) to the blotting pads and allow the system to equilibrate while to the conditions set in step 2 (∼15 minutes).
4. Using a plasma cleaner (Harrick Plasma Inc.), or equivalent system, glow discharge R2/1 200 mesh, copper grids (Quantifoil).
5. Use liquid nitrogen (LN_2_) to cool the Vitrobot foam dewar, ethane cup, and metal spider.
6. Once the setup has cooled, condense the ethane in the ethane cup. Be sure to monitor ethane and LN_2_ levels throughout the vitrification process.
7. On the Vitrobot, set the wait time to 60 seconds and set the drain time to 0.5 seconds. For blot force and blot time it is usually necessary to test a range of parameters that work best. For these fibrils a blot time between 4 and 5 seconds, and a blot force of -1 to +2 worked well.
8. Using the Vitrobot tweezers, pick up a grid and attached the tweezers to the Vitrobot. Select *“continue”* on the screen to raise the tweezers and mount the foam dewar in place. Follow the prompts on the screen to bring the tweezers and dewar into position for sample application.
9. Apply 4 μL of the fibrils to the carbon side of the grid. Select *“continue”* to begin the wait time, then the system will automatically blot and plunge the sample into liquid ethane.
10. Once the system has plunged the specimen into the cryogen, transfer the vitrified grid to a labeled grid box and store appropriately.
11. Repeat steps 7 to 9 for any additional grids. In addition to duplicate grids, it is always beneficial to test a range of blotting conditions and/or sample concentrations. Cryo-EM data was collected on a grid with a blot time of 4 seconds and a blot force of +2 at a protein concentration of ∼6.5 mg/ml.

### D. Cryo-EM data collection

Data collection parameters should be tailored to the resources available and thus, users should work closely with EM facility staff to optimize the data collection parameters for their individual sample. Here, the data was acquired on a Titan Krios G3i FEG-TEM (Thermo Fisher Scientific). The microscope is operated at 300 kV and is equipped with a Gatan K3 direct electron detector (Gatan) and a BioQuantum energy filter set at 20eV (Gatan). Correlated-double sampling (CDS) was used to collect dose fractionated micrographs using a defocus range of -0.5 to -2.5 μm with increments of 0.25 μm. Micrographs were collected at a magnification of 105,000X with a pixel size of 0.834 Å and a total dose of 40 e^-^/Å^2^ (1 e^-^/Å^2^/frame). On average ∼250 movies were collected per hour using EPU/AFIS (Thermo Fisher Scientific) acquiring 3 shots per hole and multiple holes per stage movement. A representative micrograph at an estimated defocus of -2.0 μm shows twisting fibrils suspended in vitreous ice (Figure 4).

**Figure 4.**
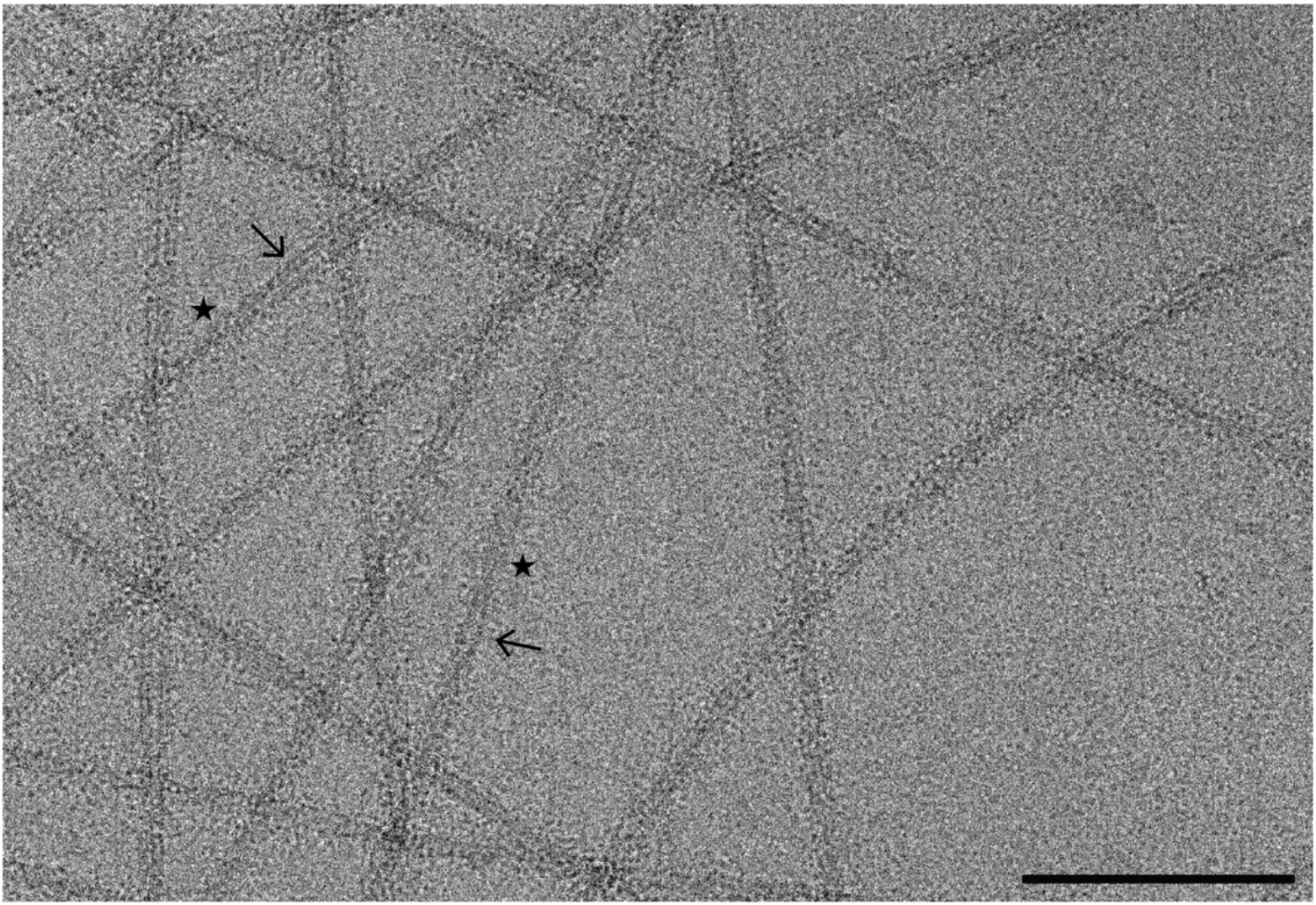
Representative cryo-EM micrographs of α-synuclein fibrils. Motion corrected micrograph of vitrified α-synuclein fibrils at an estimated defocus of -2.0 μm. The fibrils are comprised of two protofilaments (arrows) that are twisting at distinct crossover points (stars). Twisting fibrils are critical for high-resolution structure determination. Scale bar, 100 nm.

### E. Cryo-EM data processing of alpha-synuclein fibrils

Cryo-EM structure determination of amyloid fibrils has revolutionized the fields of neuroscience and neurodegenerative medicine, providing key structural details that were previously unattainable by other methods. Here, we provide a data processing protocol, that is both detailed and reproducible, to serve as a starting point for those new to cryo-EM and helical reconstruction workflows. The raw micrographs, gain file, and the detector mtf file can be accessed at EMPIAR-12229, allowing users to work through the steps below before applying the workflow to new experimental data. We must note that all data sets are unique and possess their own challenges, but this workflow should greatly improve the user’s ability to resolve amyloid fibril structures. Finally, as with any software, it is best to first become accustomed to the program by completing the appropriate tutorial datasets. We highly encourage readers to first complete the RELION single particle tutorial (https://relion.readthedocs.io/en/release-4.0/SPA_tutorial/index.html) before proceeding with the steps below [35].

<u>Creating a RELION Project</u>

Create a directory that will house the entire RELION project. For simplicity call this directory *a-syn_data_processing*. Within this directory you should have two files titled *gain*.*mrc* and *k3-CDS-300keV-mtf*.*star*, and a subdirectory titled *Micrographs* that contains all the raw movie frames in tiff format. These files can be downloaded from EMPIAR-12229. Now that your directories are organized *cd* to the *a-syn_data_processing* directory, this will serve as the RELION parent directory for all subsequent jobs. Launch RELION by running *relion &* in the terminal. The “*&*” will allow RELION to run in the background in case the terminal is needed for additional commands. As a final note, we have listed the input files for each job based on our RELION project so there will be discrepancies in job numbers between our project and yours. Thus, it is important to use the proper input path file for your project at each step. For each step we have detailed where the input file comes from (i.e. the step the file was generated in) to ensure successful reconstruction of the EMPIAR-12229 dataset.

<u>Allocating computational resource when running RELION jobs.</u>

RELION uses a *Compute* and *Running* tab to allocate computational resources based on user defined parameters. These parameters are completely dependent on the resources available to each individual. Thus, rather than detailing these parameters for each step we have outlined the *Compute* and *Running* parameters that work well for our HPC cluster with slurm queueing system here. However, these parameters may not work for your computational setup, and you may need to seek the advice of IT professionals at your institute.

<u>Compute:</u>

*Use parallel disk I/O? Yes*

*Number of pooled particles: 30*

*Skip padding? No*

*Pre-read all particles into RAM? No*

*Copy particles to scratch directory: Leave Blank*

*Combine iterations through disc? No*

*Use GPU acceleration? Yes*

*Which GPUs to use: Leave Blank*

<u>Running (GPU jobs):</u>

*Number of MPI procs: 5*

*Number of threads: 6 Submit to queue? Yes*

*Queue name: a5000*

*Queue submit command: sbatch*

*Standard submission script:* ..*/*..*/*..*/*..*/*..*/*..*/share/sbatch/relion_template_gpu*.*sh*

*Minimum dedicated cores per node: 1*

*Additional arguments: Leave Blank*

<u>Running (CPU jobs):</u>

*Number of MPI procs: 20*

*Submit to queue? Yes*

*Queue name: cpu*

*Queue submit command: sbatch*

*Standard submission script:* ..*/*..*/*..*/*..*/*..*/*..*/share/sbatch/relion_template_cpu*.*sh*

*Minimum dedicated cores per node: 1*

*Additional arguments: Leave Blank*

In addition to the protocol below, a workflow diagram of the RELION GUI with parameters for each step are provided (Figure 5).

**Figure 5.**
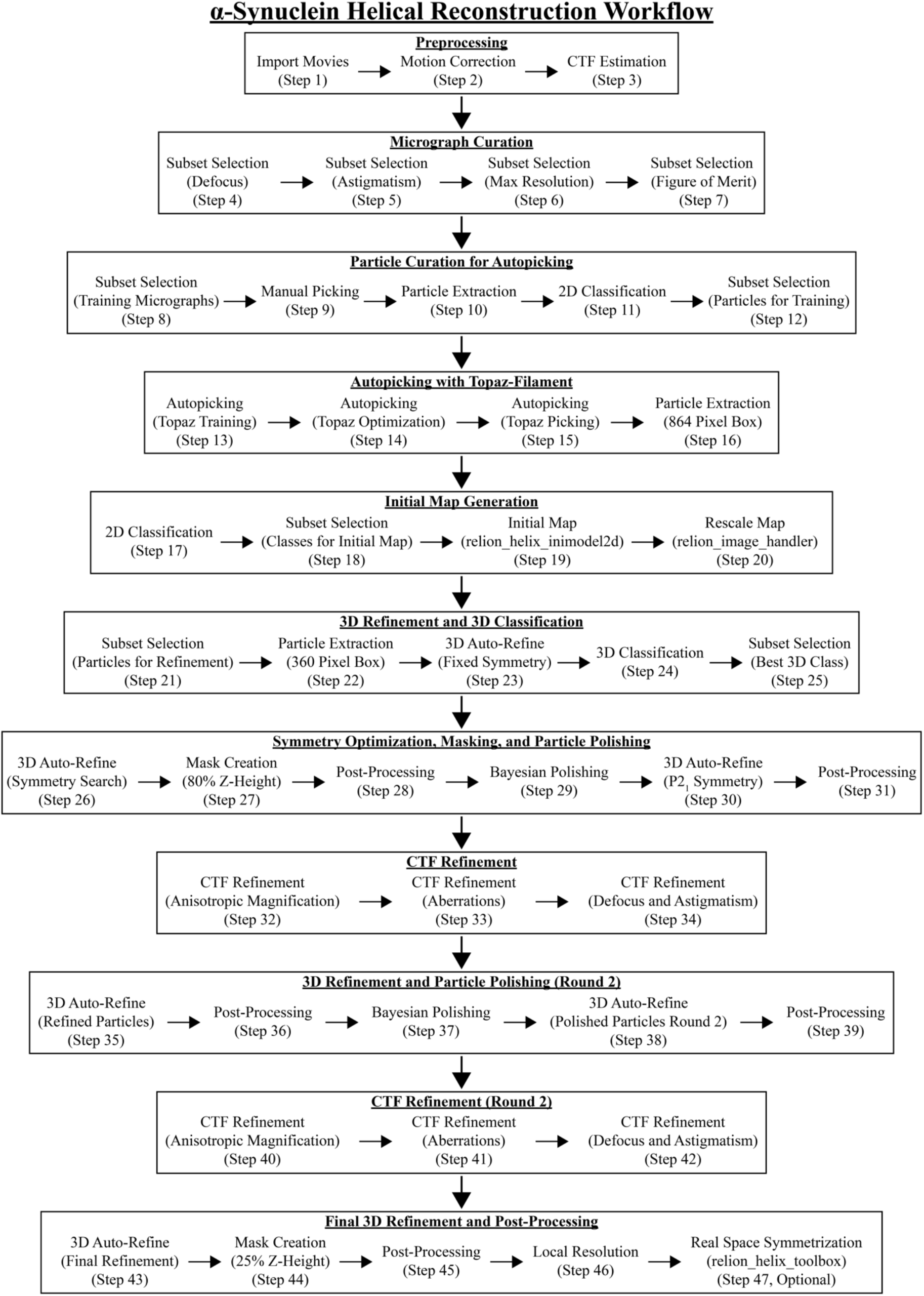
Helical reconstruction workflow for α-synuclein fibrils using RELION. Overview of each RELION job utilized to reconstruct α-synuclein fibrils to ∼2.0 Å. Each job corresponds to the step number in section E.

1. Import: First, import the raw movie frames into RELION for data processing. Select the **Import** job, ensure *Raw input files* is set to the location of the movie frames, use the “*\**” argument to select all the tiff files in the directory, set the additional parameters below, and click the *Run!* button. <u>Movies/mics:</u> *Import raw movies/micrographs:Yes* *Raw input files: Micrographs/\**.*tiff* *Optics group name: opticsGroup1* *MTF of the detector: k3-CDS-300keV-mtf*.*star* *Pixel size (Angstrom): 0*.*834* *Voltage (kV): 300* *Spherical aberration (mm): 2*.*7* *Amplitude contrast: 0*.*1* *Beamtilt in X (mrad): 0* *Beamtilt in Y (mrad): 0* <u>Others:</u> *Import other node types? No* The output log will display 5,193 micrographs were imported.
2. Motion Correction: The raw movie frames from the previous job (*movies*.*star*) must now be aligned. The data was collected with a dose per frame of 1 e^-^/Å^2^ over 40 frames for a total dose of 40 e^-^/Å^2^. Note, that the *EER fractionation* parameter will be ignored by RELION since these images were collected on a Gatan K3 detector and are tiff files. Perform motion correction by using the MotionCor2 program [36]. Tell RELION where the program is located via the *MOTIONCOR2 executable* parameter. Your computational setup will be different and MotionCor2 may be saved in a different location, so the executable path may be different. In the terminal, run *which motioncor2* to determine the correct path for the program. Similarly, your computational setup will dictate the number of GPUs available. Our setup includes multiple nodes that can each run 4 GPUs concurrently. In the RELION GUI, use *Which GPUs to use* to indicate the GPUs available for your setup, leaving this blank will automatically allocate the GPUs. Select the **Motion correction** job, set *Input movies STAR file* to the *movie*.*star* file from step 1, set the following parameters and update any paths or parameters that are specific to your computational setup, then click the *Run!* button. <u>I/O:</u> *Input movies STAR file: Import/job001/movies*.*star* *First frame for corrected sum: 1* *Last frame for corrected sum: -1* *Dose per frame (e*^*-*^*/Å*^*2*^*): 1* *Pre-exposure (e*^*-*^*/Å*^*2*^*): 0* *EER fractionation: 32 Write output in float16? Yes* *Do dose-weighting? Yes* *Save non-dose weighted as well? No* *Save sum of power spectra? Yes* *Sum power spectra every (e*^*-*^*/Å*^*2*^*): 4* <u>Motion:</u> *Bfactor: 150* *Number of patches X, Y: 5, 5* *Group frames: 1* *Binning factor: 1* *Gain-reference image: gain*.*mrc* *Gain rotation: 180 degrees (2)* *Gain flip: Flip left to right (2)* *Defect file: Leave blank* *Use RELION’s own implementation? No* *MOTIONCOR2 executable: /programs/x86_64-linux/motioncor2/1*.*3*.*1/motioncor2* *Which GPUs to use: 0,1,2,3* *Other MOTIONCOR2 arguments: Leave blank* This job will take several hours to run and will generate a *corrected_micrographs*.*star* file. If interested, you may open the *logfile*.*pdf* to visualize the results from the job. Under the *Finished Jobs* list click on the **Motion correction** job. This will update the *Current:* job display (located in the center of the GUI) to your **Motion correction** job and upload the results to the user interface. On the right side of the RELION GUI there is a drop-down menu called *Display:* that allows for the user to visualize outputs from the finished job. Click on the drop-down menu and select *Out: logfile*.*pdf*. A new window will appear with the results of the job. Subsequent *logfile*.*pdf* files from finished jobs can be opened this way.
3. CTF Estimation: Now estimate the CTF values for the motion corrected micrographs from the previous step, these are stored in the *corrected_micrographs*.*star* file. Use Gctf to estimate CTF values [37]. In the terminal, run *which Gctf* to determine the correct executable path for your setup. In the RELION GUI, select the **CTF estimation** job, set *Input micrographs STAR file* to the *corrected_micrographs*.*star* file from step 2, set the following parameters and update any paths specific to your setup, then click the *Run!* button. <u>I/O:</u> *Input micrographs STAR file: MotionCorr/job002/corrected_micrographs*.*star* *Use micrograph without dose-weighting? No* *Estimate phase shifts? No Amount of astigmatism (Å): 100* <u>CTFFIND-4.1:</u> *Use CTFFIND-4*.*1? No* *FFT box size (pix): 512* *Minimum resolution (Å): 30* *Maximum resolution (Å): 5* *Minimum defocus value (Å): 5000* *Maximum defocus value (Å): 50000* *Defocus step size (Å): 500* <u>Gctf</u> *Use Gctf instead? Yes* *Gctf executable: /programs/x86_64-linux/gctf/1*.*06/bin/Gctf* *Ignore ‘Searches’ parameters? Yes* *Perform equi-phase averaging? Yes* *Other Gctf options: Leave blank* *Which GPUs to use: 0,1,2,3* The job results in a *micrographs_ctf*.*star* file and a *logfile*.*pdf* file. The *logfile*.*pdf* contains a graphical representation of the metadata related to micrograph defocus, astigmatism, max resolution, and figure of merit values. These values will be used in the upcoming steps to filter the micrograph dataset.
4. Subset Selection (Defocus Filter): Extensive testing has shown that using stringent parameters during the micrograph curation steps allow for a segment picking neural network that performs better than one trained on the entire data set. The following steps will use CTF estimation results to curate a set of micrographs for manual picking. Those picks will then be used to train the Topaz neural network [38]. Finally, a modified version of Topaz called Topaz-filament, that allows for picking filamentous structures, is optimized on a small subset of micrographs before applying the neural network to our entire dataset [38]. Open the *logfile*.*pdf* from the **CTF estimation** job and use the values provided in this file to eliminate any outliers or suboptimal micrographs. Filter the dataset based on defocus, astigmatism, max resolution, and figure of merit values using a series of **Subset Selection** jobs. Select the **Subset Selection** job, input the following parameters and update *OR select from micrograph*.*star* to the *micrographs_ctf*.*star* file from step 3, then click the *Run!* button. <u>I/O:</u> *Select classes from job: Leave blank* *OR select from micrograph*.*star: CtfFind/job003/micrographs_ctf*.*star* *OR select from particles*.*star: Leave blank* <u>Class options:</u> *Automatically select 2D classes? No* *Re-center the class averages? No* *Regroup the particles? No* <u>Subsets:</u> *Select based on metadata value? Yes* *Metadata label for subset selection: rlnDefocusU* *Minimum metadata value: -9999* *Maximum metadata value: 25000* *OR: select on image statistics? No* *OR: split into subsets? No* <u>Duplicates:</u> *OR: remove duplicates? No* This job reduces the number of micrographs from 5,193 to 4,858 based on a maximum defocus value of 2.5 μm (25,000 Å).
5. Subset Selection (Astigmatism Filter): Filter the micrograph subset from step 4 by the astigmatism values in the **CTF estimation** *logfile*.*pdf* file. Select the **Subset Selection** job type, set *OR select from micrograph*.*star* to the *micrographs*.*star* file from step 4, input the following parameters, then click the *Run!* button. <u>I/O:</u> *Select classes from job: Leave blank* *OR select from micrograph*.*star: Select/job004/micrographs*.*star* *OR select from particles*.*star: Leave blank* <u>Class options:</u> *Automatically select 2D classes? No* *Re-center the class averages?* *No Regroup the particles? No* <u>Subsets:</u> *Select based on metadata value? Yes* *Metadata label for subset selection: rlnCtfAstigmatism* *Minimum metadata value: -9999* *Maximum metadata value: 700* *OR: select on image statistics? No* *OR: split into subsets? No* <u>Duplicates:</u> *OR: remove duplicates? No* This job reduces the number of micrographs from 4,858 to 3,390 micrographs.
6. Subset Selection (Max Resolution Filter) Further filter the micrograph subset from step 5 by the max resolution values from the **CTF estimation** *logfile*.*pdf* file. Select the **Subset Selection** job, set *OR select from micrograph*.*star* to the *micrograph*.*star* file from step 5, set the following parameters, and click the *Run!* button. <u>I/O:</u> *Select classes from job: Leave blank* *OR select from micrograph*.*star: Select/job020/micrographs*.*star* *OR select from particles*.*star: Leave blank* <u>Class options:</u> *Automatically select 2D classes? No Re-center the class averages? No Regroup the particles? No* <u>Subsets:</u> *Select based on metadata value? Yes* *Metadata label for subset selection: rlnCtfMaxResolution* *Minimum metadata value: -9999* *Maximum metadata value: 4* *OR: select on image statistics? No* *OR: split into subsets? No* <u>Duplicates:</u> *OR: remove duplicates? No* This job reduces the number of micrographs from 3,390 to 910 micrographs.
7. Subset Selection (Figure of Merit Filter): Lastly, filter the micrograph subset from step 6 by the figure of merit values from the **CTF estimation** *logfile*.*pdf* file. Select the **Subset Selection** job, set *OR select from micrograph*.*star* to the *micrograph*.*star* file from step 6, set the following parameters, then click the *Run!* button. <u>I/O:</u> *Select classes from job: Leave blank* *OR select from micrograph*.*star: Select/job021/micrographs*.*star* *OR select from particles*.*star: Leave blank* <u>Class options:</u> *Automatically select 2D classes? No* *Re-center the class averages? No* *Regroup the particles? No* <u>Subsets:</u> *Select based on metadata value? Yes* *Metadata label for subset selection: rlnCtfFigureOfMerit* *Minimum metadata value: 0*.*065* *Maximum metadata value: 0*.*9* *OR: select on image statistics? No* *OR: split into subsets? No* <u>Duplicates:</u> *OR: remove duplicates? No* This job reduces the number of micrographs from 910 to 774 micrographs.
8. Subset Selection (2 Sets of 20 Micrograph) From the remaining 774 micrographs generate 2 sets of 20 micrographs. The first set of micrographs will be used for manual picking and training the neural network. The 2^nd^ set of 20 micrographs will be used to test and optimize the picking thresholds that will then be applied to the entire dataset. Select the **Subset Selection** job, set *OR select from micrograph*.*star* to the *micrographs*.*star* file from step 7, then click the *Run!* button. <u>I/O:</u> *Select classes from job: Leave blank* *OR select from micrograph*.*star: Select/job022/micrographs*.*star* *OR select from particles*.*star: Leave blank* <u>Class options:</u> *Automatically select 2D classes? No* *Re-center the class averages? No* *Regroup the particles? No* <u>Subsets:</u> *Select based on metadata value? No* *OR: select on image statistics? No* *OR: split into subsets? Yes* *Randomise order before making subset?* *No Subset size: 20* *OR: number of subsets: 2* <u>Duplicates:</u> *OR: remove duplicates? No* This step results in two STAR files labeled *micrographs_split1*.*star* and *micrographs_split2*.*star*. Each star file contains 20 micrographs.
9. Manual Picking: Select the **Manual picking** job, set *Input micrographs* to the *micrographs_split1*.*star* file from step 8, set the additional parameters below, then click the *Run!* button. A new window will appear with 20 rows (1 per micrograph) with the micrograph name, a *pick* button, the number of picks, a *CTF* button, and the defocus estimate for that micrograph. Click on the *pick* button to launch a new window for the specified micrograph. Use the left mouse button and click at one end of a fibril, then click a 2^nd^ time at the opposite end of the fibril. This creates a line segment between the two end points defined by the user. The segments will be used for the particle extraction job in subsequent steps. Repeat this process until all the fibrils are picked. Ensure segments do not overlap, and if fibrils contain curvature increase the number of segments that make up the filament (Figure 6A). When done picking, right click on the micrograph and select *Save STAR with coordinates*, close the micrograph and repeat the process for the remaining 19 micrographs. If you need to remove points, use the center button and click over an existing point to remove it. Ensure that all the micrographs have an even number of picks (i.e. one start point and one end point per segment) and that segments are centered over fibrils. When done picking from all 20 micrographs close the window to finalize the job. <u>I/O:</u> *Input micrographs: Select/job023/micrographs_split1*.*star* *Pick start-end coordinates helices? Yes* *Use autopick FOM threshold? No* <u>Display:</u> *Particle diameter (Å): 100* *Scale for micrographs: 0*.*2* *Sigma contrast: 3* *White value: 0* *Black value: 0* *Lowpass filter (Å): 20* *Highpass filter (Å): -1* *Pixel size (Å): -1* *OR: use Topaz denoising? No* <u>Colors:</u> *Blue<>red color particles? No* The output log will list the total number of picks (start and end points). Here, we picked 414 particles (i.e. 207 segments) from 20 micrographs and the coordinates are saved to the *manualpick*.*star* file located in the directory for this job. The total number of segments may vary due to differences in picking but ensure picks are made on all 20 micrographs. NOTE: The parameters in the *Display* tab are for visualization purposes only and do not impact downstream processing steps. NOTE: We observed that in some versions of RELION there is a bug that results in an empty coordinate file from the **Manual picking** job. To bypass this error, simply select the **Manual picking** job from the *Finished jobs* section and then click on the *Continue!* button. This will reopen the manual picking GUI then, close the window; the coordinate file should now be updated with all the picks saved. There is no need to repick particles or change any settings.
10. Particle Extraction (Manual Picks): The manually picked segments must now be processed to extract particles for 2D classification. In principle, this step will take user defined parameters to then cut the segments into individual particles for downstream steps (Figure 6B). This is achieved by providing the *number of unique asymmetrical units* and *helical rise (Å)* values in the helix tab. RELION will use these values to establish an interbox distance, i.e. the spacing between each particle, that will separate overlapping 360-pixel boxes that traverse the length of the segment (Figure 6B). Here, we have set the interbox distance to ∼38.5 Å (4.82 Å x 8) to increase the number of particles for training purposes. This value will be expanded later once auto-picking is complete. Select the **Particle extraction** job, set *Micrograph STAR file* to the *micrograph_split1*.*star* file from step 8, set *Input coordinates* to the *manualpick*.*star* file from step 9, then click the *Run!* button. <u>I/O:</u> *Micrograph STAR file: Select/job023/micrographs_split1*.*star* *Input coordinates: ManualPick/job024/manualpick*.*star* *OR re-extract refined particles? No* *OR re-center refined coordinates? No* *Write output in float16? Yes* <u>Extract:</u> *Particle box size (pix): 360* *Invert contrast? Yes Normalize particles? Yes* *Diameter background circle (pix): -1* *Stddev for white dust removal: -1* *Stddev for black dust removal: -1* *Rescale particles? No* *Use autopick FOM threshold? No* <u>Helix:</u> *Extract helical segments? Yes* *Tube diameter (Å): 140* *Use bimodal angular priors? Yes* *Coordinates are start-end only? Yes* *Cut helical tubes into segments? Yes* *Number of unique asymmetrical units: 8* *Helical rise (Å): 4*.*82* This job resulted in 5,919 particles extracted to a pixel size of 0.834 Å/pix with a box size of 360 pixels. Differences in particle counts are due to differences in the number of segments picked during the manual picking step. Aim for at least 4,000 particles at this stage. NOTE: Amyloid structures have a consistent helical rise of ∼4.8 Å. This estimate is sufficient for this stage of processing, as the helical rise will be optimized in subsequent steps.
11. 2D Classification (Manual Picks): Although the particles were manually picked, and thus should be free from suboptimal picks or background noise, we prefer to perform a round of 2D classification to curate the particles that will be used to train the Topaz neural network. Select the **2D classification** job, set *Input images STAR file* to the *particles*.*star* file from step 10, set the parameters below, then click on the *Run!* button. <u>I/O:</u> *Input images STAR file: Extract/job029/particles*.*star* <u>CTF:</u> *Do CTF-correction? Yes* *Ignore CTFs until first peak? No* <u>Optimisation:</u> *Number of classes: 20* *Regularisation parameter T: 2* *Use EM algorithm? Yes* *Number of EM iterations: 20* *Use VDAM algorithm? No* *Mask diameter (Å): 285* *Mask individual particles with zeros? Yes* *Limit resolution E-step to (Å): 10* *Center class averages? Yes* <u>Sampling:</u> *Perform image alignment? Yes* *In-plane angular sampling: 2* *Offset search range (pix): 5* *Offset search step (pix): 1 Allow coarser sampling? No* <u>Helix:</u> *Classify 2D helical segments? Yes* *Tube diameter (Å): 140* *Do bimodal angular searches? Yes* *Angular search range-psi (deg): 6* *Restrict helical offsets to rise: Yes Helical rise (Å): 4*.*82* Due to the small number of particles and the small number of classes this job should only take a couple of minutes to run. The final classes can be visualized by clicking on the *Display:* drop-down menu and selecting *out: run_it020_optimiser*.*star*. A RELION display GUI will appear, check the box next to *Sort images on:* and select *rlnClassDistribution* from the drop-down menu, then click *Display!* to see the classes sorted with the most populated classes at the top (Figure 6C). Close the window when done.
12. Subset Selection (2D Classes for Topaz Training): Next, use the **Subset selection** job to select the best classes to train the Topaz neural network. Set *Select classes from job* to the *run_it020_optimiser*.*star* file from step 11, set the additional parameters below, and click the *Run!* button. This will launch a RELION display GUI. Check the box next to *Sort images on:* and select *rlnClassDisribution*, then click the *Display!* button. This will look identical to the previous step where we visualized the classes, but now use the left mouse button to select all the classes to move to the next step (Figure 6C). Once done, right click and select *Save selected classes*, then close the display window. <u>I/O:</u> *Select classes from job: Class2D/job030/run_it020_optimiser*.*star* *OR select from micrograph*.*star: Leave blank* *OR select from particles*.*star: Leave blank* <u>Class options:</u> *Automatically select 2D classes? No* *Re-center the class averages? Yes* *Regroup the particles? No* <u>Subsets:</u> *Select based on metadata values? No* *OR: select on image statistics? No* *OR: split into subsets? No* <u>Duplicates:</u> *OR: remove duplicates? No* This job resulted in 15 classes selected with 5,712 particles (Figure 6C, green boxes). Your values may be slightly different at this step due to differences in manual picking, but the key is to select classes that appear fibrillar in nature (Figure 6C, green boxes).
13. Auto-Picking (Topaz Training): Use the curated particle stack to train a new Topaz neural network. It is critical that the executable path within the *Topaz* tab directs RELION to the *topaz-filament* program [38]. The path here is to where *topaz-filament* is located on our HPC cluster, but this may be different for your setup. If you are unsure where this program is located, you may attempt to locate the program path by running the *which topaz-filament* command from the terminal. Select the **Auto-picking** job, set *Input micrographs for autopick* to the *micrographs_selected*.*star* file from step 9, in the *Topaz* tab set *Particles STAR file for training* to the *particles*.*star* file from step 12, set the additional parameters below and modify the executable path to fit your computational setup, then click on the *Run!* button. <u>I/O:</u> *Input micrographs for autopick: ManualPick/job024/micrographs_selected*.*star* *Pixel size in micrographs (Å): -1* *Use reference-based template-matching? No* *OR: use Laplacian-of-Gaussian? No* *OR: use Topaz? Yes* <u>Laplacian:</u> This tab is ignored since we opted to use Topaz in the I/O tab. <u>Topaz:</u> *Topaz executable: /programs/x86_64-linux/system/sbgrid_bin/topaz-filament* *Particle diameter (Å): 140* *Perform topaz picking? No* *Perform topaz training? Yes* *Nr of particle per micrograph: 300* *Input picked coordinates for training: Leave blank* *OR train on a set of particles? Yes* *Particles STAR file for training: Select/job032/particles*.*star* *Additional topaz arguments: Leave blank* <u>References:</u> This tab is ignored since we opted to use Topaz in the I/O tab. <u>Autopicking:</u> *Use GPU acceleration? Yes* All other parameters on this tab are ignored since we opted to use Topaz in the I/O tab. <u>Helix:</u> This tab is ignored since we opted to use Topaz in the I/O tab. This job results in a trained Topaz model titled *model_epoch10*.*sav* and is saved in the folder for this job. NOTE: Topaz training is not parallelized so the job will only use 1 MPI process.
14. Auto-Picking (Topaz Picking Optimization) The trained Topaz model will be applied to a subset of 20 micrographs to test how the model performs before it is applied to the entire dataset. For *topaz-filament* to pick segments, and not individual particles as in traditional single particle analysis, the additional flags for filament (*-f)* and threshold (-t) must be provided in the *Additional topaz arguments* box. Additionally, an integer value must be provided after the threshold flag. This threshold determines how many particles are picked. A lower threshold results in more particles, but if the threshold is too low, then the model will start picking noise. With any new trained Topaz neural network, we test a range of threshold values, typically from -6 to 0, to see which threshold works best (Figure 6D). Each threshold value will be its own job. Select the **Auto-picking** job, set *Input micrographs for autopick* to the *micrographs_split2* from job 8, in the *Topaz* tab set *Trained topaz model* to the *model_epoch10*.*sav* file from step 13, set the parameters below, and click the *“Run!”* button. To test additional thresholds, once the first **Auto-picking** job is complete, click on the job in the *Finished jobs* list and then click on the **Auto-picking** job to load the previous settings. Now, simply change the threshold value in the *Additional topaz arguments* and click the *“Run!” button*. Repeat this process for any threshold that you would like to test. We tested thresholds -6, -5, -4, -3, -2, -1, and 0, and found that threshold -5 worked best for the dataset (Figure 6D). We have also included an extreme case with a threshold of -10 to better visualize bad picks that would be unsuitable for further processing (Figure 6D). <u>I/O:</u> *Input micrographs for autopick: Select/job023/micrographs_split2*.*star* *Pixel size in micrographs (Å): -1* *Use reference-based template-matching? No* *OR: use Laplacian-of-Gaussian? No* *OR: use Topaz? Yes* <u>Laplacian:</u> This tab is ignored since we opted to use Topaz in the I/O tab. <u>Topaz:</u> *Topaz executable: /programs/x86_64-linux/system/sbgrid_bin/topaz-filament* *Particle diameter (Å): 140* *Perform topaz picking? Yes* *Trained topaz model: AutoPick/job033/model_epoch10*.*sav* *Perform topaz training? No* *Additional topaz arguments: -f -t -5* <u>References:</u> This tab is ignored since we opted to use Topaz in the I/O tab. <u>Autopicking:</u> *Use GPU acceleration? Yes* All other parameters on this tab are ignored since we opted to use Topaz in the I/O tab. <u>Helix:</u> This tab is ignored since we opted to use Topaz in the I/O tab. A picking threshold of -5 resulted in 688 segments (1,376 particles, i.e. end points) from 20 micrographs. NOTE: Topaz picking is parallelized so multiple MPI processes can be run simultaneously, we typically run 20 MPI processes for this job. This setting can be found in the *Running* tab and is dependent on the computational resources available.
15. Auto-Picking (Topaz Picking on the Entire Dataset) The trained Topaz model and the optimized picking threshold are now applied to the entire dataset to select segments for downstream processing. As detailed previously, upload the settings from the best picking job (threshold -5), update *Input micrographs for autopick* to the *micrographs*.*star* file from step 4, and click the *“Run!”* button. <u>I/O:</u> *Input micrographs for autopick: Select/job004/micrographs*.*star* *Pixel size in micrographs (Å): -1* *Use reference-based template-matching? No* *OR: use Laplacian-of-Gaussian? No* *OR: use Topaz? Yes* <u>Laplacian:</u> This tab is ignored since we opted to use Topaz in the I/O tab. <u>Topaz:</u> *Topaz executable: /programs/x86_64-linux/system/sbgrid_bin/topaz-filament* *Particle diameter (Å): 140* *Perform topaz picking? Yes* *Trained topaz model: AutoPick/job033/model_epoch10*.*sav* *Perform topaz training? No* *Additional topaz arguments: -f -t -5* <u>References:</u> This tab is ignored since we opted to use Topaz in the I/O tab. <u>Autopicking:</u> *Use GPU acceleration? Yes* All other parameters on this tab are ignored since we opted to use Topaz in the I/O tab. <u>Helix:</u> This tab is ignored since we opted to use Topaz in the I/O tab. This job results in 156,526 segments (313,052 particles, i.e. end points) from 4,858 micrographs.
16. Particle Extraction (Large Box Size): For helical reconstruction methods, the helical twist and rise values are critical for cryo-EM data processing. The helical twist can be estimated from 2D class averages with large box sizes that span the fibril crossover distance (Figure 7A, 7B, 7D). Here, extract the particles to a box size of 864 pixels (∼720 Å) so we can estimate the crossover distance in subsequent steps. At this stage in the processing there is no need for high resolution information, so the box size is rescaled to 144 pixels (i.e. binning to a pixel size of 5.004 Å/pixel). Alternatively, users may estimate the crossover distance from cryo-EM micrographs, typically those with higher defocus values are easier to visualize, or from negative stain TEM micrographs. However, extraction at a larger box size is still necessary to generate an initial reference for 3D reconstruction. Select the **Particle extraction** job, set *Micrograph STAR file* to the *micrographs*.*star* file from step 4, set *Input coordinates* to the *autopick*.*star* file from step 15, set the additional parameters below, and click the *“Run!”* button. <u>I/O:</u> *Micrograph STAR file: Select/job004/micrographs*.*star* *Input coordinates: AutoPick/job041/autopick*.*star* *OR re-extract refined particles? No* *OR re-center refined coordinates? No* *Write output in float16? Yes* <u>Extract:</u> *Particle box size (pix): 864* *Invert contrast? Yes* *Normalize particles? Yes* *Diameter background circle (pix): -1* *Stddev for white dust removal: -1* *Stddev for black dust removal: -1* *Rescale particles? Yes* *Re-scale size (pixels): 144* *Use autopick FOM threshold? No* <u>Helix:</u> *Extract helical segments? Yes* *Tube diameter (Å): 140* *Use bimodal angular priors? Yes* *Coordinates are start-end only? Yes* *Cut helical tubes into segments? Yes* *Number of unique asymmetrical units: 15* *Helical rise (Å): 4*.*82* This job results in 771,754 particles with an original box size of 864 pixels that is rescaled to 144 pixels at a pixel size of 5.004 Å/pixel. NOTE: The number of asymmetrical units was increased to 15, this results in an interbox distance of ∼72 Å or ∼25% of the small box size (360 pixels) that will be used for the final reconstruction.
17. 2D Classification (Large Box Size) Classify the particles to remove junk particles and to estimate the crossover distance. Select the **2D classification** job, set *Input images STAR file* to the *particles*.*star* file from step 16, set the additional parameters below, then click on the “Run!” button. <u>I/O:</u> *Input images STAR file: Extract/job042/particles*.*star* <u>CTF:</u> *Do CTF-correction? Yes* *Ignore CTFs until first peak? Yes* <u>Optimisation:</u> *Number of classes: 50* *Regularisation parameter T: 2* *Use EM algorithm? Yes* *Number of EM iterations: 20* *Use VDAM algorithm? No* *Mask diameter (Å): 710* *Mask individual particles with zeros? Yes* *Limit resolution E-step to (Å): -1* *Center class averages? Yes* <u>Sampling:</u> *Perform image alignment? Yes* *In-plane angular sampling: 2* *Offset search range (pix): 5* *Offset search step (pix): 1* *Allow coarser sampling? No* <u>Helix:</u> *Classify 2D helical segments? Yes* *Tube diameter (Å): 140* *Do bimodal angular searches? Yes* *Angular search range-psi (deg): 6* *Restrict helical offsets to rise: Yes* *Helical rise (Å): 4*.*82* This job results in the *run_it020_optimiser*.*star* file that contains the 2D class averages. This file can be viewed using the *Display:* drop-down menu on the right side of the GUI. NOTE: Sometimes it can be helpful to determine the helical rise of the filament rather than assume 4.8 Å as the starting point. To do this, users can utilize 2D classifications and use measurements of the average power spectra in Fourier space to calculate the estimated rise. To perform this analysis, use a box size of 360 pixels and high-resolution data (0.834 Å/pix), as this allows for more detail to be visualized in the 2D classes (specifically the β-sheet rungs). To do so, use the **Particle Extraction** job to extract particles to their original pixel size. Use the parameters as instructed in step 16, but ensure that *Particle box size* is set to *360* and that *Rescale particles* is set to *No*. Once particle extraction is complete, run a **2D Classification** job as described in step 17. Ensure *Input images STAR files* is set to the correct *particles*.*star* file from the **Particle Extraction** job, and *Mask diameter* is set to *300*. When the job is done, open the average power spectra by selecting the *out:run_it020_optimiser* from the display output. Enter an increased *Sigma Contrast* value in the top box of the RELION display GIU (we used 1 for our data) (Figure 7C). If the user fails to increase the Sigma Contrast, the average power spectra will not be visible (Figure 7C). Once the 2D classes are displayed, right click on a class and select *Show Fourier amplitudes (2X)*. This will open an image of the average power spectra. Make a measurement from the meridian to either layer line with the strong intensity (Figure 7C). This can be done by clicking and holding the center button on the mouse. Use the following formula to calculate the rise: *rise* 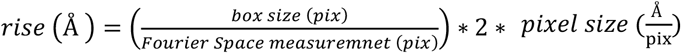 (Figure 7C, 7D). For new experimental data, if the rise is substantially different then parameters for steps 16 onward should reflect the updated rise. Here, a measurement of ∼124 pixels results in a helical rise of 4.84 Å that will be refined in later steps (Figure 7C).
18. Subset Selection (2D Classes for Initial Map) Select 2-3 good classes that will be used to generate an initial 3D volume. Select the **Subset Select** job, set *Select classes from job* to the *run_it020_optimiser*.*star* file from step 17, set the parameters below, then click the *Run!* button. A RELION display GUI will appear, reverse sort the class averages by *rlnClassDistribution* (as described in step 11) and select 2 class averages (Figure 8A, green boxes). To measure the crossover distance right click on a 2D class and select *Show original image*. A new window will appear, using the center button click and drag to measure the distance between two crossovers (Figure 7B). The distance in pixels is displayed over the image and in the terminal (Figure 7B). Multiply the measured distance by the current pixel size of 5.004 Å/pixel to calculate the distance in angstroms (Figure 7D). Here, we estimated a crossover distance of 120 pixels or 600 Å (Figure 7B). When done, close the original image. Repeat the process for any additional 2D classes you want to measure. Lastly, in the window with all the 2D classes, right click and select *Save STAR with selected images*, then close display window. <u>I/O:</u> *Select classes from job: Class2D/job044/run_it020_optimiser*.*star* *OR select from micrograph*.*star: Leave blank* *OR select from particles*.*star: Leave blank* <u>Class options:</u> *Automatically select 2D classes? No* *Re-center the class averages? Yes* *Regroup the particles? No* <u>Subsets:</u> *Select based on metadata values? No* *OR: select on image statistics? No* *OR: split into subsets? No* <u>Duplicates:</u> *OR: remove duplicates? No* This job results in a *class_averages*.*star* file containing the 2 selected classes.
19. Initial Map Generation Using *relion_helix_inimodel2d* Generate an initial map from the selected 2D classes in step 18 using the *relion_helix_inimodel2d* program [39]. The following steps must be completed in the terminal. First create two directories to keep our data organized. In the terminal, navigate to the RELION project directory, this is the directory that contains all the RELION subdirectories, and enter the commands *mkdir inimodel* and *mkdir ini4refine*. This will create two directories one that will house the initial volumes and a second that will contain the rescaled volumes that will be used for refinement steps. Documentation on *relion_helix_inimodel2d* can be found at https://relion.readthedocs.io/en/release-4.0/Reference/Helix.html. For convenience, we have detailed each argument below, alternatively running *relion_helix_inimodel2d* with no additional arguments will detail all available arguments for the program. Before running the command, ensure that the input STAR file (*--i)* and the output root name (*--o)* are updated to your specific project, then run the command from the terminal. The following command generates an initial volume with an estimated crossover distance of 750 Å (Figure 8F). *relion_helix_inimodel2d --i Select/job045/class_averages*.*star --angpix 5*.*004 --mask_diameter 300 --sym 2* *--iter 10 --search_shift 70 --search_angle 15 --search_size 10 --j 20 --crossover_distance 750 --o inimodel/Select045_CO750* The arguments run with *relion_helix_inimodel2d* are detailed below. *--i* input STAR file with 2D classes *--angpix* pixel size in angstroms *--mask_diameter* size in angstroms of circular mask around 2D classes *--sym* order of symmetry in 2D slices *--iter* number of iterations to run *--search_shift* distance in angstroms to search translations perpendicular to helical axis *--search_angle* degrees to search in-plane rotations *--search_size* ± number of pixels to fix best crossover distance *--j* number of threads *--crossover_distance* distance in angstroms between 2 crossovers *--o* output root name The program generates several files, and the initial 3D volume is saved with the suffix *_class001_rec3d*.*mrc* which can be opened in ChimeraX for visualization [40-42]. Since the initial crossover distance is an estimate, we prefer to generate several initial maps for a round of 3D refinement to see what best fits our experimental data (Figure 8B-8G). Although our initial estimate for crossover distance was 600 Å, we found that an initial map with a crossover distance of 750 Å is best for this dataset (Figure 8C, 8F). You may test additional crossover distances as we typically do with new experimental datasets. To do so, change the *--crossover_distance* and *--o* arguments of the command above to generate additional maps of varying crossover distances with appropriate output root names (Figure 8B-8G).
20. Rescale Initial Map Using *relion_image_handler* The initial maps generated in the previous step must be rescaled because the 3D refinement steps will be performed with a smaller box size (360 pixels) at the original pixel size (0.834 Å/pixel). Use *relion_image_handler* to rescale the maps. For a list of all possible arguments simply run the program in the terminal with no additional arguments. The command below was used to rescale the 750 Å crossover map. Before running the command ensure the MRC input file (*--i)* and the MRC output file (*--o)* reflect your project. For convenience, the arguments used to run *relion_image_handler* are detailed below. *relion_image_handler --i inimodel/Select045_CO750_class001_rec3d*.*mrc --angpix 5*.*004 --rescale_angpix 0*.*834 --new_box 360 --o ini4refine/Select045_CO750_box360*.*mrc* *--i* input MRC file of the initial map *--angpix* pixel size in angstroms of the input file *--rescale_angpix* scale input map to this new pixel size in angstroms *--new_box* resize the input map to this box size in pixels *--o* output name of resized map Repeat this step for any additional maps that will be tested. Ensure that the input file (*--i)* and output file (*--o*) are updated to reflect the maps being rescaled (Figure 8B-8G).
21. Subset Selection (2D Classes for Refinement) Select additional classes from the **2D Classification** job (step 17) to ensure there are enough particles for additional processing. Repeat the **Subset Selection** job as in step 18 but now select all the good classes for further processing (Figure 8A). <u>I/O:</u> *Select classes from job: Class2D/job044/run_it020_optimiser*.*star* *OR select from micrograph*.*star: Leave blank* *OR select from particles*.*star: Leave blank* <u>Class options:</u> *Automatically select 2D classes? No* *Re-center the class averages? Yes* *Regroup the particles? No* <u>Subsets:</u> *Select based on metadata values? No* *OR: select on image statistics? No* *OR: split into subsets? No* <u>Duplicates:</u> *OR: remove duplicates? No* Here, we selected 21 classes with 413,249 particles saved in the *particles*.*star* file (Figure 8A).
22. Particle Extraction (Small Box Size): Select the **Particle Extraction** job, set *Refined particles STAR files* to the *particles*.*star* file from step 21, set the additional parameters below, and click the *Run!* button. <u>I/O:</u> *Micrograph STAR file: Leave blank* *Input coordinates: Leave blank* *OR re-extract refined particles? Yes* *Refined particles STAR file: Select/job046/particles*.*star* *Reset the refined offsets to zero? Yes* *OR re-center refined coordinates? No* *Write output in float16? Yes* <u>Extract:</u> *Particle box size (pix): 360* *Invert contrast? Yes* *Normalize particles? Yes* *Diameter background circle (pix): -1* *Stddev for white dust removal: -1* *Stddev for black dust removal: -1* *Rescale particles? No* *Use autopick FOM threshold? No* <u>Helix:</u> *Extract helical segments? Yes* *Tube diameter (Å): 140* *Use bimodal angular priors? Yes* *Coordinates are start-end only? Yes* *Cut helical tubes into segments? Yes* *Number of unique asymmetrical units: 15* *Helical rise (Å): 4*.*82* The 413,249 particles were re-extracted to a box size of 360 pixels and a pixel size of 0.834 Å/pixel. The particles are stored in the *particles*.*star* file.
23. 3D Auto-Refine (Fixed Symmetry) The particle set from step 22 and the rescaled initial map generated in step 20 will be subjected to a round of 3D refinement. First, take the estimated helical rise and calculate the initial twist for the estimated crossover distance using the following formula: 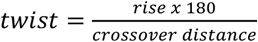 (Figure 7D). The rise is estimated to be 4.82 Å and the crossover distance was estimated to be 750 Å, so the initial twist value is 1.16°. Finally, apply a negative value to the initial twist based on the assumption that fibrils typically display a left-handed helical form and as supported by atomic force microscopy studies [13]. Select the **3D Auto-Refine** job, set *Input images STAR file* to the *particles*.*star* file generated in step 22, set *Reference map* to the rescaled initial volume generated in step 20 in our case this file was named *Select045_CO750_box360*.*mrc*, set the parameters below, and click the *Run!* button. <u>I/O:</u> *Input images STAR file: Extract/job047/particles*.*star* *Reference map: ini4refine/Select045_CO750_box360*.*mrc* *Reference mask (optional): Leave blank* <u>Reference:</u> *Ref. map is on absolute greyscale? No Initial low-pass filter (Å): 10 Symmetry: C1* <u>CTF:</u> *Do CTF-correction? Yes* *Ignore CTFs until first peak? No* <u>Optimisation:</u> *Mask diameter (Å): 220* *Mask individual particles with zeros? Yes* *Use solvent-flattened FSCs? No* <u>Auto-sampling:</u> *Initial angular sampling: 3*.*7 degrees* *Initial offset range (pix): 5* *Initial offset step (pix): 1* *Local searches from auto-sampling: 1*.*8 degrees* *Relax symmetry: Leave blank* *Use finer angular sampling faster? (No)* <u>Helix:</u> *Do helical reconstruction? Yes* *Tube diameter – inner, outer (Å): -1, 140* *Angular search range – rot, tilt, psi (deg): -1, 15, 10* *Range factor of local averaging: -1* *Keep tilt-prior fixed: Yes* *Apply helical symmetry? Yes* *Number of unique asymmetrical units: 15* *Initial twist (deg), rise (Å): -1*.*16, 4*.*82* *Central Z length (%): 25* *Do local searches of symmetry? No* The job results in a map with a global resolution of 3.66 Å (Figure 9A, 9B, blue). Repeat this step for any additional initial maps and crossover distances that you would like to test. We tested crossover distances of 550, 600, 650, 700, 750, and 800 Å (Figure 9A, 9B). These tests showed that three maps resolved to a resolution of 3.66 Å. The map generated from the 750 Å crossover distance was selected because the backbone density was best resolved, and the map showed side chain densities for some residues (Figure 9B, blue). Additionally, the map showed clear separation of the β-strands along the helical axis.
24. 3D Classification (Symmetry Search) During the **Subset Selection** job (step 21), we selected all the 2D classes that resembled amyloid fibrils. Being less stringent after 2D classification means that heterogeneity most likely exists in our dataset. By using 3D classification, we can further sort the heterogeneity that may exist in the particle set and improve the quality of the reconstruction. Use the 3D reconstruction from step 23 as an initial starting point to then sort particles into 4 classes. Use the *Do local searches of symmetry* tool to search a range of helical parameters that best fit the dataset. Select the **3D Classification** job, set *Input images STAR file* to the *run_data*.*star* file from step 23, set *Reference map* to the *run_half1_class001_unfil*.*mrc* from step 23, set the additional parameters below, and click the *Run!* button. <u>I/O:</u> *Input images STAR file: Refine3D/job069/run_data*.*star* *Reference map: Refine3D/job069/run_half1_class001_unfil*.*mrc* *Reference mask (optional): Leave blank* <u>Reference:</u> *Ref. map is on absolute greyscale? No* *Initial low-pass filter (Å): 4*.*5* *Symmetry: C1* <u>CTF:</u> *Do CTF-correction? Yes* *Ignore CTFs until first peak? No* <u>Optimization:</u> *Number of classes: 4* *Regularization parameter T: 4* *Number of iterations: 20* *Use fast subsets (for large data sets)? No* *Mask diameter (Å): 220* *Mask individual particles with zeros? Yes* *Limit resolution E-step to (Å): -1* <u>Sampling:</u> *Perform image alignment? Yes* *Angular sampling interval: 3*.*7 degrees* *Offset search range (pix): 5* *Offset search step (pix): 1* *Perform local angular searches? No* *Allow coarser sampling? No* <u>Helix:</u> *Do helical reconstruction? Yes* *Tube diameter – inner, outer (Å): -1, 140* *Angular search range – rot, tilt, psi (deg): -1, 15, 10* *Range factor of local averaging: -1* *Keep tilt-prior fixed: Yes* *Apply helical symmetry? Yes* *Number of unique asymmetrical units: 15* *Initial twist (deg), rise (Å): -1*.*14, 4*.*82* *Central Z length (%): 25* *Do local searches of symmetry? Yes* *Twist search – Min, Max, Step (deg): -0*.*9, -1*.*2, 0*.*01* *Rise search – Min, Max, Step (Å): 4*.*75, 4*.*95, 0*.*01* The job runs for 20 iterations, sorting the particle set into 4 classes and optimizing helical parameters at each iteration. A cross section of the 3D volumes can be visualized by displaying the *run_it020_optimiser*.*star* file in RELION. Alternatively, the 4 MRC files generated in this job (*run_it020_class001*.*mrc, run_it020_class002*.*mrc*, etc.) can be opened in ChimeraX for easier visualization of the 3D maps. Class 3 was the best 3D volume with a helical twist of -1.12° and a helical rise of 4.84 Å (Figure 9C, green box).
25. Subset Selection (3D Class for Additional Processing) Use the **Subset selection** job to select the best class from the 3D classification job in step 24. Ensure *Select classes from job* is set to the *run_it020_optimiser*.*star* file that was generated in step 24. Set the parameters below and click the *Run!* button. Refer to step 12 for how to display, select, and save classes in a **Subset selection** job. <u>I/O:</u> *Select classes from job: Class3D/job077/run_it020_optimiser*.*star* *OR select from micrograph*.*star: Leave blank* *OR select from particles*.*star: Leave blank* <u>Class options:</u> *Automatically select 2D classes? No* *Re-center the class averages? Yes* *Regroup the particles? No* <u>Subsets:</u> *Select based on metadata values? No* *OR: select on image statistics? No* *OR: split into subsets? No* <u>Duplicates:</u> *OR: remove duplicates? No* Class 3 was selected in this job and the data was saved to the *particles*.*star* file that contained all 129,940 particles for that class (Figure 9C).
26. 3D Auto-Refine (Symmetry Search) Select the **3D auto-refine** job, update *Input images STAR file* to the *particles*.*star* file from step 25 and the *Reference map* to best 3D map from the 3D classification in step 24, in our case this was *run_it020_class003*.*mrc*, but this may be different for your project. Then set the parameters below and click the *Run!* button. <u>I/O:</u> *Input images STAR file: Select/job080/particles*.*star* *Reference map: Class3D/job077/run_it020_class003*.*mrc* *Reference mask (optional): Leave blank* <u>Reference:</u> *Ref. map is on absolute greyscale? No* *Initial low-pass filter (Å): 4*.*5* *Symmetry: C1* <u>CTF:</u> *Do CTF-correction? Yes* *Ignore CTFs until first peak? No* <u>Optimisation:</u> *Mask diameter (Å): 220* *Mask individual particles with zeros? Yes* *Use solvent-flattened FSCs? No* <u>Auto-sampling:</u> *Initial angular sampling: 3*.*7 degrees* *Initial offset range (pix): 5* *Initial offset step (pix): 1* *Local searches from auto-sampling: 1*.*8 degrees* *Relax symmetry: Leave blank* *Use finer angular sampling faster? (No)* <u>Helix:</u> *Do helical reconstruction? Yes* *Tube diameter – inner, outer (Å): -1, 140* *Angular search range – rot, tilt, psi (deg): -1, 15, 10* *Range factor of local averaging: -1* *Keep tilt-prior fixed: Yes* *Apply helical symmetry? Yes* *Number of unique asymmetrical units: 15* *Initial twist (deg), rise (Å): -1*.*11, 4*.*84* *Central Z length (%): 25* *Do local searches of symmetry? Yes* *Twist search – Min, Max, Step (deg): -0*.*9, -1*.*3, 0*.*01* *Rise search – Min, Max, Step (Å): 4*.*75, 4*.*95, 0*.*01* The optimized helical parameters converged to a helical twist of -1.11° and a rise of 4.84 Å. The resolution without masking is 3.23 Å. The *run_class001*.*mrc* file can be downloaded and opened with ChimeraX to visual the 3D volume (Figure 10, step 26).
27. Mask Creation (80% Mask) Helical reconstruction is prone to loss of resolvability as the volume reaches the edge of the box. Thus, masking encompasses a central portion of the fibril and excludes the ends of the fibril. The mask can be as small the as the *Central Z length* established in the **3D auto-refine** job. However, at this stage in processing we may benefit from a larger mask to ensure we have sufficient signal for the CTF refinement steps. Open the *run_class001*.*mrc* file from step 26 in ChimeraX. Ensure that the volume step is set to 1 then lower the volume threshold until noise starts to appear in the solvent space. Note this threshold and set this as the *Initial binarization threshold* for the **Mask creation** job. A value of 0.00096 worked well for this project. Update the *Input 3D map* to the *run_class001*.*mrc* generated in the step 26. Set the additional parameters below then click the *Run!* button. <u>I/O:</u> *Input 3D map: Refine3D/job081/run_class001*.*mrc* <u>Mask:</u> *Lowpass filter map (Å) 15* *Pixel size (Å) -1* *Initial binarization threshold: 0*.*00096* *Extend binary map this many pixels: 5* *Add a soft-edge of this many pixels: 5* <u>Helix:</u> *Mask a 3D helix? Yes* *Central Z length (%): 80* In ChimeraX, open the *mask*.*mrc* file and the *run_class001*.*mrc* file from step 26. Ensure both maps are set to a step size of 1, set the mask threshold to 0.99 to visualize the mask volume, and for easier visualization lower the mask opacity to 50% (Figure 11A). Inspect the mask and map, when viewing the central cross-section of the map ensure the entire proteinaceous volume is within the mask. If there are no issues, then proceed to the next step. However, if the map is not completely encompassed by the mask, lower the *Initial binarization threshold* value and rerun the job by clicking the *Continue!* button. Repeat this process until the mask is satisfactory (Figure 11A).
28. Post-Processing The post-processing job will recalculate the global resolution with masking, and it will automatically estimate and apply a B-factor to sharpen the map, further improving the quality of the map. Select the **Post-processing** job, set *One of the 2 unfiltered half-maps* to the *run_half1_class001_unfil*.*mrc* file from step 26, set *Solvent mask* to the *mask*.*mrc* file from step 27, set *MTF of the detector (STAR file)* to the *k3-CDS-300keV-mtf*.*star* file that is supplied with EMPIAR-12229. Set the remaining parameters below then click the *Run!* button. <u>I/O:</u> *One of the 2 unfiltered half-maps: Refine3D/job081/run_half1_class001_unfil*.*mrc* *Solvent mask: MaskCreate/job086/mask*.*mrc* *Calibrated pixel size (Å) -1* <u>Sharpen:</u> *Estimate B-factor automatically? Yes* *Lowest resolution for auto-B fit (Å): 10* *Use your own B-factor? No* *Skip FSC-weighting? No* *MTF of the detector (STAR file): k3-CDS-300keV-mtf*.*star* *Original detector pixel size: -1* The job estimated a b-factor of -97, and the processed map is saved as *postprocess*.*mr*c. The job also calculated a resolution of 2.97 Å with masking and the volume is saved as *postprocess_mask*.*mrc* (Figure 10, step 28).
29. Bayesian Polishing (Round 1) The next steps will aim at improving the quality of the particles to further improve the resolvability of the map. The polishing will use motion corrected micrographs and particle positions to improve motion correction on a per-particle basis. Select the **Bayesian polishing** job, set the *Micrographs (from MotionCorr)* to the *corrected_micrographs*.*star* file from step 2, set the *Particles (from Refine 3D or CtfRefine)* to the *run_data*.*star* file from step 26, set the *Postprocess STAR file* to the *postprocess*.*star* file from step 28, set the remaining parameters below, and click the *Run!* button. <u>I/O:</u> *Micrographs (from MotionCorr): MotionCorr/job002/corrected_micrographs*.*star* *Particles (from Refine 3D or CtfRefine): Refine3D/job081/run_data*.*star* *Postprocess STAR file: PostProcess/job088/postprocess*.*star* *First movie frame: 1* *Last movie frame: -1* *Extraction size (pix in unbinned movie): -1* *Re-scale size (pixels): -1* *Write output in float16? Yes* <u>Train:</u> *Train optimal parameters? No* <u>Polish:</u> *Perform particle polishing? Yes* *Optimized parameter file: Leave blank* *OR use your own parameters?* *Sigma for velocity (Å/dose): 0*.*2* *Sigma for divergence (Å): 5000* *Sigma for acceleration (Å/dose): 2* *Minimum resolution for B-factor fit (Å): 20* *Maximum resolution for B-factor fit (Å): -1* The job will save the particles to the *shiny*.*star* file. The improvements of the particle positions can be found in the *logfile*.*pdf* file.
30. 3D Auto-Refine (Pseudo-Screw Symmetry) Up to this point we have only applied helical symmetry to the 3D reconstruction. We will now address additional symmetry that may be present to further improve the quality of the reconstruction. Previous studies have shown that amyloid fibrils exist with varying degrees of symmetry. For two protofilament fibrils, we observe either C2 symmetry, where two protofilaments are identical and in register, as commonly observed in Tau fibrils, or we observe pseudo-screw symmetry (P2_1_), where two protofilaments are identical but out of register, as observed in α-synuclein fibrils (Figure 1G, 1F) [12,13,43]. To understand this difference in symmetry, it is necessary to manually inspect the reconstruction to determine the best symmetry for the dataset. This can be done by using ChimeraX to analyze the 3D volume from either the *run_class001*.*mrc* file from step 26 or the *postprocess*.*mrc* file from step 28. Here, we determined that pseudo-screw symmetry exists within our dataset. To apply this symmetry, we will continue to set the *Symmetry* parameter to C1, but we will divide the helical rise in half and subtract the helical twist from 180°. By doing so, we can impose pseudo-screw symmetry to our reconstruction. Select the **3D auto-refine** job, set *Input images STAR files* to the *shiny*.*star* file generated in step 29, set *Reference map* to the *run_half1_class001_unfil*.*mrc* file from job 26, set the additional parameters below, then click the *Run!* button. <u>I/O:</u> *Input images STAR file: Polish/job091/shiny*.*star* *Reference map: Refine3D/job081/run_half1_class001_unfil*.*mrc* *Reference mask (optional): Leave blank* <u>Reference:</u> *Ref. map is on absolute greyscale? No* *Initial low-pass filter (Å): 4*.*5* *Symmetry: C1* <u>CTF:</u> *Do CTF-correction? Yes* *Ignore CTFs until first peak? No* <u>Optimization:</u> *Mask diameter (Å): 220* *Mask individual particles with zeros? Yes* *Use solvent-flattened FSCs? No* <u>Auto-sampling:</u> *Initial angular sampling: 3*.*7 degrees* *Initial offset range (pix): 5* *Initial offset step (pix): 1* *Local searches from auto-sampling: 1*.*8 degrees* *Relax symmetry: Leave blank* *Use finer angular sampling faster? (No)* <u>Helix:</u> *Do helical reconstruction? Yes* *Tube diameter – inner, outer (Å): -1, 140* *Angular search range – rot, tilt, psi (deg): -1, 15, 10* *Range factor of local averaging: -1* *Keep tilt-prior fixed: Yes* *Apply helical symmetry? Yes* *Number of unique asymmetrical units: 15* *Initial twist (deg), rise (Å): 179*.*445, 2*.*42* *Central Z length (%): 25* *Do local searches of symmetry? Yes* *Twist search – Min, Max, Step (deg): 179*.*24, 179*.*65, 0*.*01* *Rise search – Min, Max, Step (Å): 2*.*2, 2*.*6, 0*.*01* The unmasked reconstruction improved from 3.23 Å (step 26) to 3.00 Å and is stored in the *run_class001*.*mrc* file (Figure 10, step 30). The symmetry parameters reflect pseudo-screw symmetry and were optimized to a twist of 179.45° and a rise of 2.42 Å.
31. Post-Processing Run a **Post-processing** job to see how masking the solvent region improves the resolution and how automated sharpening can improve the map quality. Select the **Post-processing** job, set *One of the 2 unfiltered half-maps* to the *run_half1_class001_unfil*.*mrc* file from step 30, set the *Solvent mask* to the *mask*.*mrc* file from step 27, set the additional parameters below, then click the *Run!* button. <u>I/O:</u> *One of the 2 unfiltered half-maps: Refine3D/job093/run_half1_class001_unfil*.*mrc* *Solvent mask: MaskCreate/job086/mask*.*mrc* *Calibrated pixel size (Å) -1* <u>Sharpen:</u> *Estimate B-factor automatically? Yes* *Lowest resolution for auto-B fit (Å): 10* *Use your own B-factor? No* *Skip FSC-weighting? No* *MTF of the detector (STAR file): k3-CDS-300keV-mtf*.*star* *Original detector pixel size: -1* Use ChimeraX to visualize the improvements to the *postprocess_masked*.*mrc* map. The GS-FSC_0.143_ for this map with masking improved from 2.97 Å to 2.89 Å and sharpening improved side chain densities throughout the map (Figure 10, step 31).
32. CTF Refinement (Anisotropic Magnification, Round 1) The next three steps will utilize the **CTF refinement** job to improve the CTF fits for the particle set. The three jobs perform corrections for 1) anisotropic magnification, 2) asymmetrical and symmetrical aberrations, and 3) recalculates per-particle defocus and per-micrograph astigmatism. Together these steps improve CTF fits that translate into improvements in the reconstruction. The first job will correct for anisotropic magnification. Select the **CTF refinement** job, set the *Particles (from Refine3D)* to the *run_data*.*star* file from step 30, set the *Postprocess STAR file* to the *postprocess*.*star* file from step 31, set the parameters below, then click the *Run!* button. <u>I/O:</u> *Particles (from Refine3D): Refine3D/job093/run_data*.*star* *Postprocess STAR file: PostProcess/job095/postprocess*.*star* <u>Fit:</u> *Estimate (anisotropic) magnification? Yes* *Minimum resolution for fits (Å): 30* This job estimated a magnification anisotropy of 0.31% and stored the refined particles to the *particles_ctf_refine*.*star* file.
33. CTF Refinement (Asymmetrical and Symmetrical Aberrations, Round 1) Use the refined particles from the previous job to correct for asymmetrical and symmetrical aberrations. Select the **CTF refinement** job, set *Particles (from Refine3D)* to the *particles_ctf_refine*.*star* file from step 32, set *Postprocess STAR file* to the *postprocess*.*star* file from step 31, set the parameters below, then click the *Run!* button. <u>I/O:</u> *Particles (from Refine3D): CtfRefine/job096/particles_ctf_refine*.*star* *Postprocess STAR file: PostProcess/job095/postprocess*.*star* <u>Fit:</u> *Estimate (anisotropic) magnification? No* *Perform CTF parameter fitting? Yes* *Fit defocus? Per-particle* *Fit astigmatism? Per-micrograph* *Fit B-factor? No* *Fit phase-shift? No* *Estimate beamtilt? No* *Estimate 4*^*th*^ *order aberrations? No* *Minimum resolution for fits (Å): 30* The refined particles are stored in the *particles_ctf_refine*.*star* file and the results of the job can be visualized by opening the *logfile*.*pdf*.
34. CTF Refinement (Recalculate Defocus and Astigmatism, Round 1) Next, recalculate defocus values on a per-particle basis and astigmatism on a per-micrograph basis. Select the **CTF refinement** job, set *Particles (from Refine3D)* to the *particles_ct_refine*.*star* file from step 33, set *Postprocess STAR file* to the *postprocess*.*star* file from step 31, set the additional parameters below, then click the *Run!* button. <u>I/O:</u> *Particles (from Refine3D): CtfRefine/job097/particles_ctf_refine*.*star* *Postprocess STAR file: PostProcess/job095/postprocess*.*star* <u>Fit:</u> *Estimate (anisotropic) magnification? No* *Perform CTF parameter fitting? No* *Estimate beamtilt? Yes* *Also estimate trefoil? Yes* *Estimate 4*^*th*^ *order aberrations? Yes* *Minimum resolution for fits (Å): 30* The refined particles are saved to the *particles_ctf_refine*.*star* file and are now ready for 3D refinement.
35. 3D Auto-Refine (CTF Refined Particles, Round 1) Generate a new 3D volume with the refined particles. Select the **3D auto-refine** job, set *Input images STAR file* to the *particles_ctf_refine*.*star* file from step 34, set *Reference map* to the *run_half1_class001*.*mrc* file from step 30, set the parameters below, ensure that the helical parameters are updated to the optimized twist and rise values from step 30 (these are found in the output log from step 30), then click the *Run!* button. <u>I/O:</u> *Input images STAR file: CtfRefine/job098/particles_ctf_refine*.*star* *Reference map: Refine3D/job093/run_half1_class001_unfil*.*mrc* *Reference mask (optional): Leave blank* <u>Reference:</u> *Ref. map is on absolute greyscale? No* *Initial low-pass filter (Å): 4*.*5* *Symmetry: C1* <u>CTF:</u> *Do CTF-correction? Yes* *Ignore CTFs until first peak? No* <u>Optimization:</u> *Mask diameter (Å): 220* *Mask individual particles with zeros? Yes* *Use solvent-flattened FSCs? No* <u>Auto-sampling:</u> *Initial angular sampling: 3*.*7 degrees* *Initial offset range (pix): 5* *Initial offset step (pix): 1* *Local searches from auto-sampling: 1*.*8 degrees* *Relax symmetry: Leave blank* *Use finer angular sampling faster? (No)* <u>Helix:</u> *Do helical reconstruction? Yes* *Tube diameter – inner, outer (Å): -1, 140* *Angular search range – rot, tilt, psi (deg): -1, 15, 10* *Range factor of local averaging: -1* *Keep tilt-prior fixed: Yes* *Apply helical symmetry? Yes* *Number of unique asymmetrical units: 15* *Initial twist (deg), rise (Å): 179*.*448, 2*.*42* *Central Z length (%): 25* *Do local searches of symmetry? Yes* *Twist search – Min, Max, Step (deg): 179*.*24, 179*.*65, 0*.*01* *Rise search – Min, Max, Step (Å): 2*.*2, 2*.*6, 0*.*01* After CTF refinement, the resolution of the unmasked 3D reconstruction improved from 3.00 Å to 2.38 Å (Figure 10, step 35). The helical parameters did not change, and converged to a twist of 179.45° and a rise of 2.42 Å. The 3D map is saved to the *run_class001*.*mrc* file and can be opened in ChimeraX for visualization.
36. Post-Processing Apply the mask from step 27 to recalculate the FSC and sharpen the map. Select the **Post-processing** job, set *One of the 2 unfiltered half-maps* to the *run_half1_class001_unfil*.*mrc* file from step 35, set *Solvent mask* to the *mask*.*mrc* file from step 27, set the additional parameters below, and click the *Run!* button. <u>I/O:</u> *One of the 2 unfiltered half-maps: Refine3D/job099/run_half1_class001_unfil*.*mrc* *Solvent mask: MaskCreate/job086/mask*.*mrc* *Calibrated pixel size (Å) -1* <u>Sharpen:</u> *Estimate B-factor automatically? Yes* *Lowest resolution for auto-B fit (Å): 10* *Use your own B-factor? No* *Skip FSC-weighting? No* *MTF of the detector (STAR file): k3-CDS-300keV-mtf*.*star* *Original detector pixel size: -1* The B-factor was estimated to -55 and applied to the map. The resolution of the masked map improved from 2.89 Å to 2.31 Å (Figure 10, step 36). The 3D map was saved to the *postprocess_masked*.*mrc* file and can be visualized in ChimeraX.
37. Bayesian Polishing (Round 2) Perform one more cycle of polishing and CTF refinement before a final round of 3D refinement and postprocessing (step 29-36). Select the **Bayesian polishing** job, set *Micrographs (from MotionCorr)* to the *corrected_micorgraphs*.*star* file from step 2, set *Particles from Refine 3D or CtfRefine* to the *run_data*.*star* file from step 35, set *Postprocess STAR file* to the *postprocess*.*star* file from step 36, set the additional parameters below, and click the *Run!* button. <u>I/O:</u> *Micrographs (from MotionCorr): MotionCorr/job002/corrected_micrographs*.*star* *Particles from Refine 3D or CtfRefine: Refine3D/job099/run_data*.*star* *Postprocess STAR file: PostProcess/job100/postprocess*.*star* *First movie frame: 1* *Last movie frame: -1* *Extraction size (pix in unbinned movie): -1* *Re-scale size (pixels): -1* *Write output in float16? Yes* <u>Train:</u> *Train optimal parameters? No* <u>Polish:</u> *Perform particle polishing? Yes* *Optimized parameter file: Leave blank* *OR use your own parameters?* *Sigma for velocity (Å/dose): 0*.*2* *Sigma for divergence (Å): 5000* *Sigma for acceleration (Å/dose): 2* *Minimum resolution for B-factor fit (Å): 20* *Maximum resolution for B-factor fit (Å): -1* The polished particles are stored in the *shiny*.*star* file.
38. 3D Auto-Refine (Polished Particles, Round 2) Use the polished particles from step 37 and perform a round of 3D refinement. Select the **3D auto-refine** job, set *Input images STAR files* to the *shiny*.*star* file from step 37, set *Reference map* to the *run_half1_class001_unfil*.*mrc* file from step 35, set the parameters below, and click the *Run!* button. <u>I/O:</u> *Input images STAR file: Polish/job101/shiny*.*star* *Reference map: Refine3D/job099/run_half1_class001_unfil*.*mrc* *Reference mask (optional): Leave blank* <u>Reference:</u> *Ref. map is on absolute greyscale? No* *Initial low-pass filter (Å): 4*.*5* *Symmetry: C1* <u>CTF:</u> *Do CTF-correction? Yes* *Ignore CTFs until first peak? No* <u>Optimization:</u> *Mask diameter (Å): 220* *Mask individual particles with zeros? Yes* *Use solvent-flattened FSCs? No* <u>Auto-sampling:</u> *Initial angular sampling: 3*.*7 degrees* *Initial offset range (pix): 5* *Initial offset step (pix): 1* *Local searches from auto-sampling: 1*.*8 degrees* *Relax symmetry: Leave blank* *Use finer angular sampling faster? (No)* <u>Helix:</u> *Do helical reconstruction? Yes* *Tube diameter – inner, outer (Å): -1, 140* *Angular search range – rot, tilt, psi (deg): -1, 15, 10* *Range factor of local averaging: -1* *Keep tilt-prior fixed: Yes* *Apply helical symmetry? Yes* *Number of unique asymmetrical units: 15* *Initial twist (deg), rise (Å): 179*.*449, 2*.*42* *Central Z length (%): 25* *Do local searches of symmetry? Yes* *Twist search – Min, Max, Step (deg): 179*.*24, 179*.*65, 0*.*01* *Rise search – Min, Max, Step (Å): 2*.*2, 2*.*6, 0*.*01* After a 2^nd^ round of polishing the unmasked map did not improve in resolution, staying at 2.38 Å (Figure 10, step 38). Next, we will see if there is an improvement in the masked reconstruction.
39. Post-Processing Select the **Post-processing** job, set *One of the 2 unfiltered half-maps* to the *run_half1_class001_unfil*.*mrc* file from step 38, set the *Solvent mask* to the *mask*.*mrc* file from step 27, set the additional parameters below, and click the *Run!* button. <u>I/O:</u> *One of the 2 unfiltered half-maps: Refine3D/job102/run_half1_class001_unfil*.*mrc* *Solvent mask: MaskCreate/job086/mask*.*mrc* *Calibrated pixel size (Å) -1* <u>Sharpen:</u> *Estimate B-factor automatically? Yes* *Lowest resolution for auto-B fit (Å): 10* *Use your own B-factor? No* *Skip FSC-weighting? No* *MTF of the detector (STAR file): k3-CDS-300keV-mtf*.*star* *Original detector pixel size: -1* The resolution of the masked reconstruction increased slightly from 2.31 Å to 2.27 Å (Figure 10, step 39). The 3D map is stored in the *postprocess_masked*.*mrc* file and can be visualized in ChimeraX.
40. CTF Refinement (Anisotropic Magnification, Round 2) Perform a final round of CTF refinements as in steps 32-34. Select the **CTF refinement** job, set *Particles (from Refine3D)* to the *run_data*.*star* file from step 38, set *Postprocess STAR file* to the *postprocess*.*star* file from step 39, set the parameters below, then click the *Run!* button. <u>I/O:</u> *Particles (from Refine3D): Refine3D/job102/run_data*.*star* *Postprocess STAR file: PostProcess/job103/postprocess*.*star* <u>Fit:</u> *Estimate (anisotropic) magnification? Yes* *Minimum resolution for fits (Å): 30* The refined particles are stored in the *particles_ctf_refine*.*star* file and will be used in the next step.
41. CTF Refinement (Asymmetrical and Symmetrical Aberrations, Round 2) Select the **CTF refinement** job, set *Particles (from Refine3D)* to the *particles_ctf_refine*.*star* file from step 40, set *Postprocess STAR file* to the *postprocess*.*star* file from step 39, set the parameters below, then click the *Run!* button. <u>I/O:</u> *Particles (from Refine3D): CtfRefine/job104/particles_ctf_refine*.*star* *Postprocess STAR file: PostProcess/job103/postprocess*.*star* <u>Fit:</u> *Estimate (anisotropic) magnification? No* *Perform CTF parameter fitting? Yes* *Fit defocus? Per-particle* *Fit astigmatism? Per-micrograph* *Fit B-factor? No* *Fit phase-shift? No* *Estimate beamtilt? No* *Estimate 4*^*th*^ *order aberrations? No* *Minimum resolution for fits (Å): 30* The particles were written out to the *particles_ctf_refine*.*star* file and will be used in the next step.
42. CTF Refinement (Recalculate Defocus and Astigmatism, Round 2) Select the **CTF refinement** job, set *Particles (from Refine3D)* to the *particles_ctf_refine*.*star* file from step 41, set *Postprocess STAR files* to the *postprocess*.*star* file from step 39, set the additional parameters below, then click the *Run!* button. <u>I/O:</u> *Particles (from Refine3D): CtfRefine/job105/particles_ctf_refine*.*star* *Postprocess STAR file: PostProcess/job103/postprocess*.*star* <u>Fit:</u> *Estimate (anisotropic) magnification? No* *Perform CTF parameter fitting? No* *Estimate beamtilt? Yes* *Also estimate trefoil? Yes* *Estimate 4*^*th*^ *order aberrations? Yes* *Minimum resolution for fits (Å): 30* The refined particles are stored in the *particls_ctf_refine*.*star* file and are ready for 3D refinement.
43. 3D Auto-Refine (CTF Refined Particles, Round 2) Run a 3D refinement using the CTF refined particles. Select the **3D auto-refine** job, set *Input images STAR file* to the *particles_ctf_refine*.*star* file from step 42, set *Reference map* to the *run_half1_class001_unfil*.*mrc* file from step 38, set the additional parameters below, and click the *Run!* button. <u>I/O:</u> *Input images STAR file: CtfRefine/job106/particles_ctf_refine*.*star* *Reference map: Refine3D/job102/run_half1_class001_unfil*.*mrc* *Reference mask (optional): Leave blank* <u>Reference:</u> *Ref. map is on absolute greyscale? No* *Initial low-pass filter (Å): 4*.*5* *Symmetry: C1* <u>CTF:</u> *Do CTF-correction? Yes* *Ignore CTFs until first peak? No* <u>Optimization:</u> *Mask diameter (Å): 220* *Mask individual particles with zeros? Yes* *Use solvent-flattened FSCs? No* <u>Auto-sampling:</u> *Initial angular sampling: 3*.*7 degrees* *Initial offset range (pix): 5* *Initial offset step (pix): 1* *Local searches from auto-sampling: 1*.*8 degrees* *Relax symmetry: Leave blank* *Use finer angular sampling faster? (No)* <u>Helix:</u> *Do helical reconstruction? Yes* *Tube diameter – inner, outer (Å): -1, 140* *Angular search range – rot, tilt, psi (deg): -1, 15, 10* *Range factor of local averaging: -1* *Keep tilt-prior fixed: Yes* *Apply helical symmetry? Yes* *Number of unique asymmetrical units: 15* *Initial twist (deg), rise (Å): 179*.*449, 2*.*42* *Central Z length (%): 25* *Do local searches of symmetry? Yes* *Twist search – Min, Max, Step (deg): 179*.*24, 179*.*65, 0*.*01* *Rise search – Min, Max, Step (Å): 2*.*2, 2*.*6, 0*.*01* The resolution of the unmasked map increased slightly from 2.38 Å to 2.35 Å (Figure 10, step 43). This result suggests that any additional rounds of polishing or CTF refinement will not yield meaningful gains in map quality and are thus not necessary.
44. Mask Creation (25% Mask) In the *Helix* tab of the **3D auto-refine** job we set the *central Z length* to 25% of the particle box. This central region is where searching for helical symmetry occurs and is also the region where real-space helical symmetry is imposed. In the previous **Mask creation** job from step 27, the mask length was set to 80% of the central axis to ensure enough signal was available for CTF refinements. Now, in the final stages of processing we can reduce the mask size to a *central Z length* of 25% as was used in the 3D reconstruction steps. As in step 27, you may need to open the *run_class001*.*mrc* file from step 43 in ChimeraX to determine the appropriate *Initial binarization threshold* for the reconstruction. A value of 0.0011 worked for well for us. Select the **Mask creation** job, set *Input 3D map* to the *run_class001*.*mrc* file from step 43, set the parameters below, then click the *Run!* button. <u>I/O:</u> *Input 3D map: Refine3D/job107/run_class001*.*mrc* <u>Mask:</u> *Lowpass filter map (Å) 15* *Pixel size (Å) -1* *Initial binarization threshold: 0*.*0011* *Extend binary map this many pixels: 5* *Add a soft-edge of this many pixels: 5* <u>Helix:</u> *Mask a 3D helix? Yes* *Central Z length (%): 25* The mask is saved to the *mask*.*mrc* file and will be used in the next step (Figure 11B).
45. Post-Processing Apply the latest mask from step 44 to the final reconstruction from step 43 to recalculate the resolution and B-factor. Select the **Post-processing** job, set *One of the 2 unfiltered half-maps* to the *run_half1_class001_unfil*.*mrc* file from step 43, set *Solvent mask* to the *mask*.*mrc* file from step 44, set the additional parameters below, then click the *Run!* button. <u>I/O:</u> *One of the 2 unfiltered half-maps: Refine3D/job107/run_half1_class001_unfil*.*mrc* *Solvent mask: MaskCreate/job109/mask*.*mrc* *Calibrated pixel size (Å) -1* <u>Sharpen:</u> *Estimate B-factor automatically? Yes* *Lowest resolution for auto-B fit (Å): 10* *Use your own B-factor? No* *Skip FSC-weighting? No* *MTF of the detector (STAR file): k3-CDS-300keV-mtf*.*star* *Original detector pixel size: -1* The final masked map has a resolution of 2.04 Å and a B-factor of -40 (Figure 10, step 45). The map displays well resolved side chain densities as expected for a map at ∼2 Å resolution. NOTE: We observe a spike in the FSC plot at ∼2.4 Å (the repeating unit) in both our reconstruction and in several published structures (Figure 12) [22,27,44]. This spike is alleviated with masking, but it is a common feature observed in amyloid structures that resolve to high resolution. Additionally, other helical structures, such as tad pili, also display a similar spike due to the strong signal at the repeating unit of ∼4.9 Å [45].
46. Local resolution Calculate a local resolution map to understand the differences in resolution across the map. Select the **Local resolution** job, set *One of the 2 unfiltered half-maps* to the *run_half1_class001_unfil*.*mrc* file from step 43, set *User-provided solvent mask* to the *mask*.*mrc* file from step 44, set the additional parameters below, then click the *Run!* button. <u>I/O:</u> *One of the 2 unfiltered half-maps: Refine3D/job107/run_half1_class001_unfil*.*mrc* *User-provided solvent mask: MaskCreate/job109/mask*.*mrc* *Calibrated pixel size (Å): 0*.*834* <u>ResMap:</u> *Use ResMap? No* <u>Relion:</u> *Use Relion? Yes* *User-provided B-factor: -40* *MTF of the detector (STAR file): k3-CDS-300keV-mtf*.*star* The job results in a *histogram*.*pdf* file that contains a graph of the local resolution within the provided mask. The *relion_locres*.*mrc* file can be opened in ChimeraX along with the *postprocess*.*mrc* file from step 45 to color the surface of the map by resolution (Figure 13). Please see the “Analyzing the results” section in the RELION local resolution documentation page for details on handling these maps in ChimeraX (https://relion.readthedocs.io/en/latest/SPA_tutorial/Validation.html).
47. Real-Space Symmetrization (Optional) As stated previously, real-space symmetry is applied to only the central 25% of the reconstruction and the molecular model is built into this central region (Figure 1E). However, in some cases it is beneficial to extend the symmetrization to the edge of the box. For example, to better visualize the crossover distance we generate a map with real-space symmetry imposed to the edge of the box, then we align several models in ChimeraX to generate a multi-map volume that spans close to 1000 Å (Figure 1A). This process allows for easier visualization of the crossover distance when making figures. To impose real-space symmetry run the *relion_helix_toolbox* command in the terminal. Before running the command, *cd* to the job directory for step 45. *relion_helix_toolbox --impose --i postprocess_masked*.*mrc --o postprocess_masked_sym*.*mrc --* *cyl_outer_diameter 220 --angpix 0*.*834 --rise 2*.*42 --twist 179*.*45 --z_percentage 0*.*25* The arguments used in the command above are as follow: *--impose* apply real-space helical symmetry *--i* input file *--o* output file *--cyl_outer_diameter* outer diameter of the cylindrical mask *--angpix* pixel size in angstroms *--rise* helical rise in angstroms *--twist* helical twist in degrees *--z_percentage* central z-length

**Figure 6.**
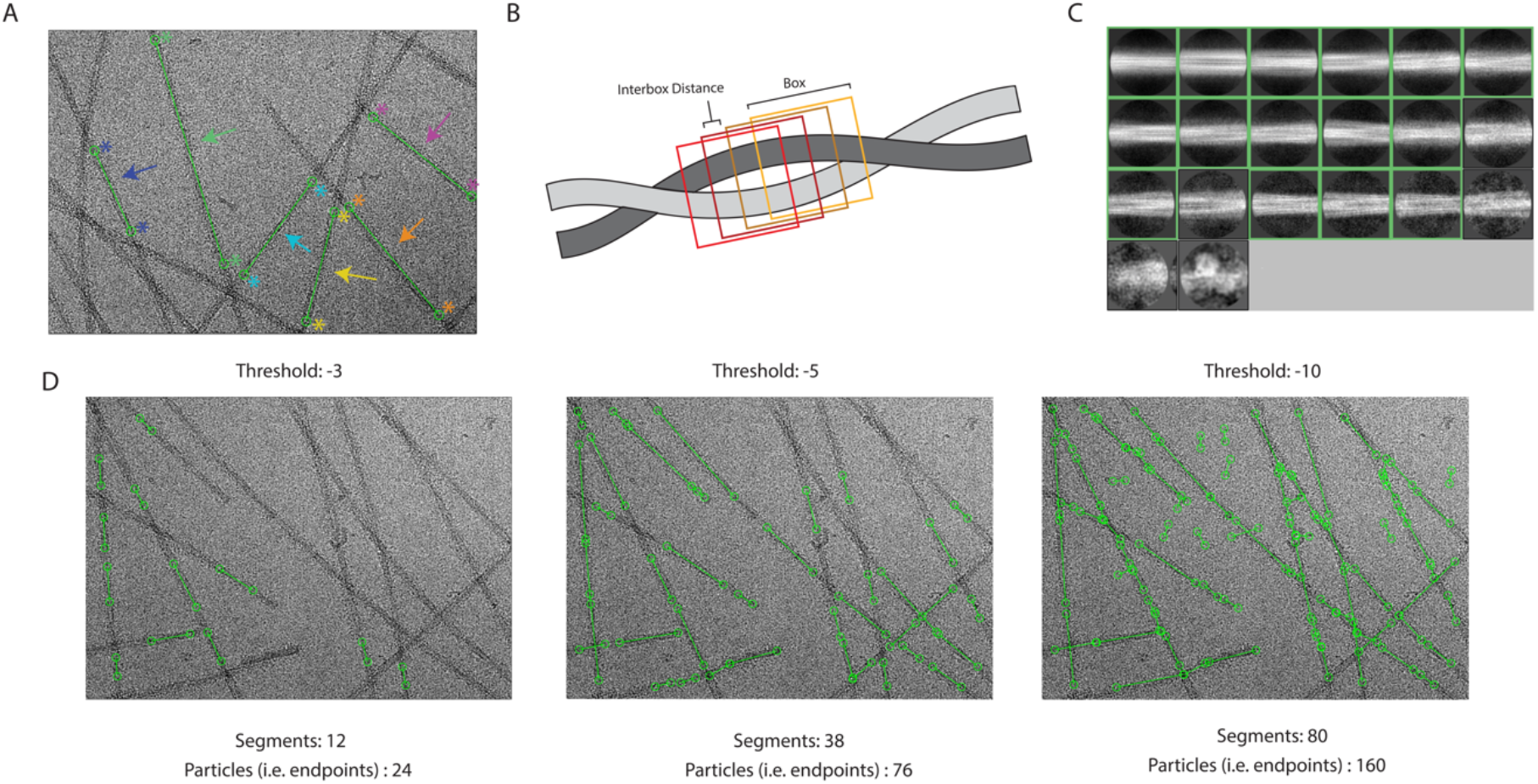
Manual picking, 2D class selection, and auto-picking threshold determination. A. Micrograph with examples of manually picked segments (step 9). Each “end” of the segment is selected by the user (indicated by the stars). The endpoints are then linked by a line (indicated by an arrow), this region will be divided into particles based on the user defined interbox distance. Each new color represents a new segment that has been manually picked. B. Schematic of interbox distances (step 10). The filament that is shown is a region that has been selected for particle picking. RELION will use a user defined “box” to select as a particle. The interbox distance shown is the distance in which no overlap from previous boxes is present (i.e., the region that is unique to each box). C. 2D Classes from manually picked particles (step 11). The green boxes indicate the classes selected to use for neural network training (step 12). D. Micrographs depicting trained neural network auto-picking results from different threshold values. As the threshold for picking is decreased, the stringency in which the neural network determines whether the feature fits the trained model is decreased—first resulting in an increase in picked particles, however, as the threshold continues to decrease, the neural network starts to categorize “noise” as pickable particles.

**Figure 7.**
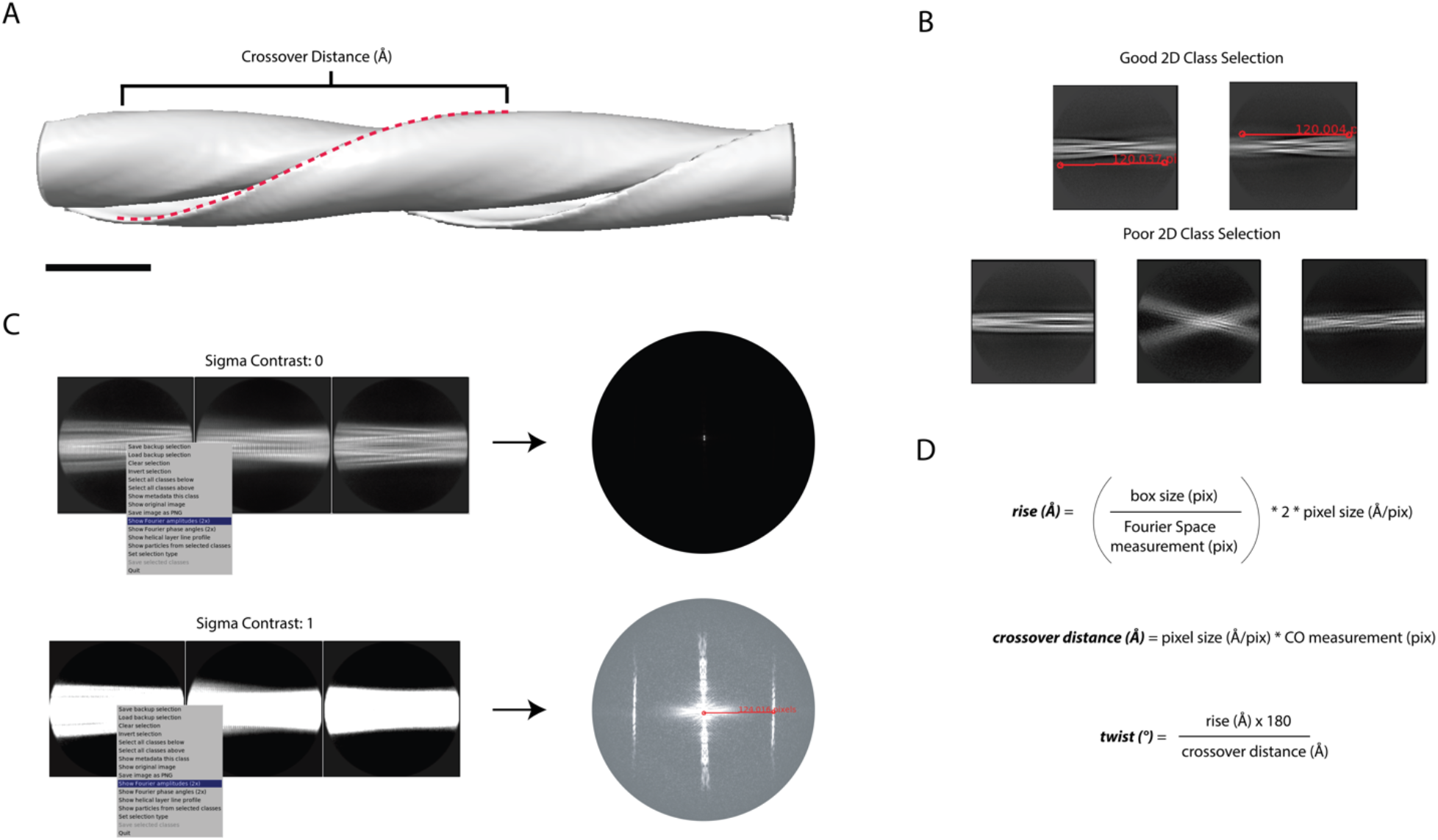
Determining crossover distance, helical twist, and helical rise. A. An initial map depicts the crossover distance observed in twisting fibrils. The crossover distance is described as the length where the fibril turns 180° (red dotted line). Scale bar, 100 nm. B. The crossover distance can be measured (red line) from well aligned 2D classes where the twisting nature of the fibril is observed, this requires a box size that spans a distance that is close to or larger than the crossover distance for an accurate measurement to be made. Here, a box size of 864 pixels (720 Å) was used for initial crossover estimates. Poor 2D classes are mis-aligned or blurry preventing crossover distance measurements. C. The helical rise can be determined from 2D classes with a small box size (360 pixels) extracted at their original pixel size (0.834 Å/pix) that yield high resolution details (i.e. spacing of the β-sheets). The sigma contrast of the 2D classes must be adjusted to visualize the helical layer lines in reciprocal space. From the average power spectrum, a measurement (red line) can be made from the meridian to the highest intensity layer line, this measurement can be used to estimate the helical rise. D. The measurements made in B and C are used to calculate the helical rise and the crossover distance. Then, the crossover distance and helical rise are used to calculate the helical twist of the structure. The estimated helical parameters are used for subsequent 3D refinement steps.

**Figure 8.**
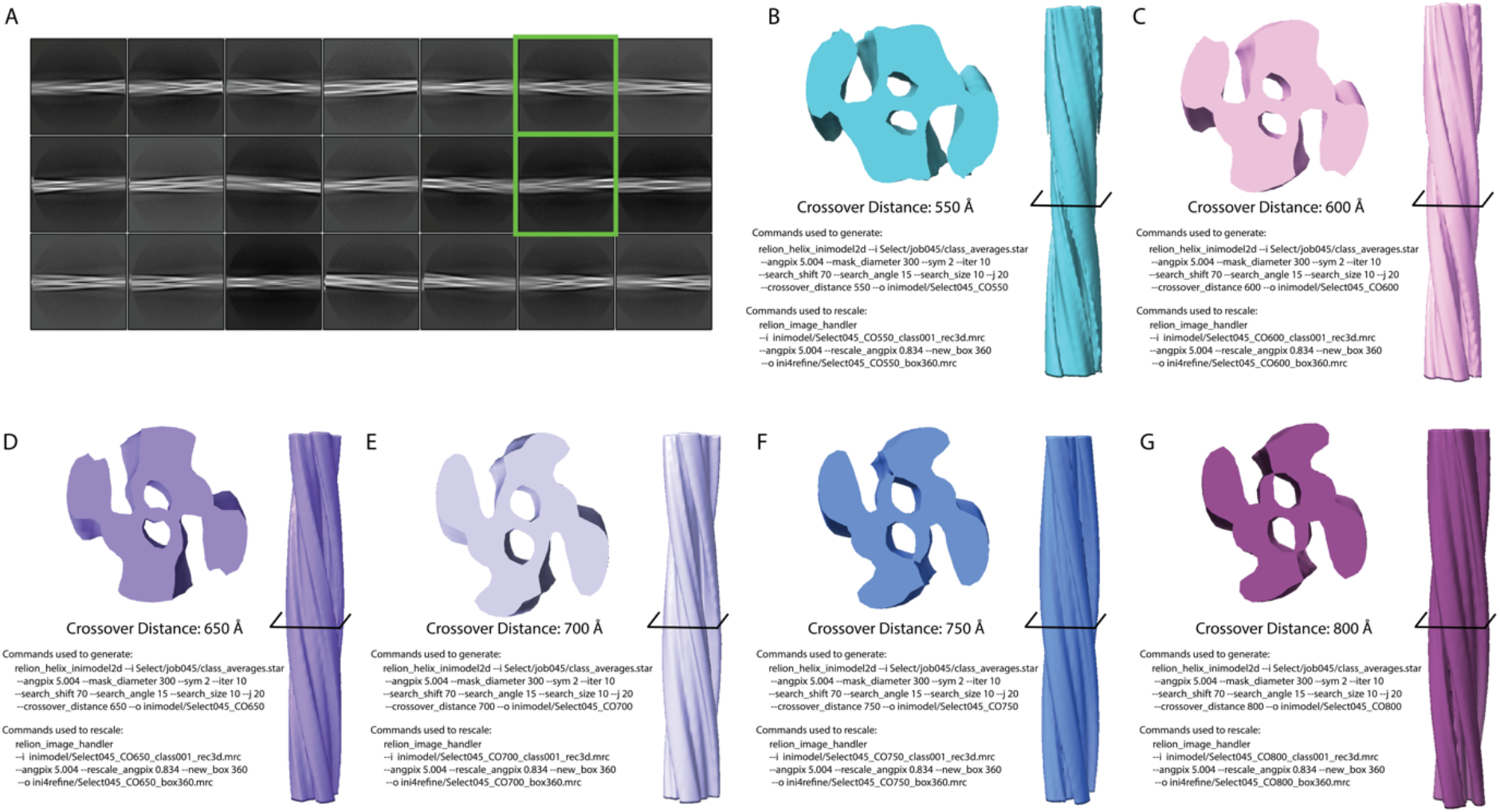
Initial model generation. A. Classes selected from all 2D classes. All classes shown in A are the classes selected (job 21) from the classes rendered from the trained neural network auto-picking job on all micrographs (step 17). The green boxes indicate the two classes selected for initial model generation (step 18). B-G. Initial maps for the crossover distances 550-800 Å. One showing the cross-section of the refined filament (cross-section location shown by the black crossbar) and the other depicting the entirety of the filament. The commands used to generate (step 19) and rescale (step 20) the initial models are shown.

**Figure 9.**
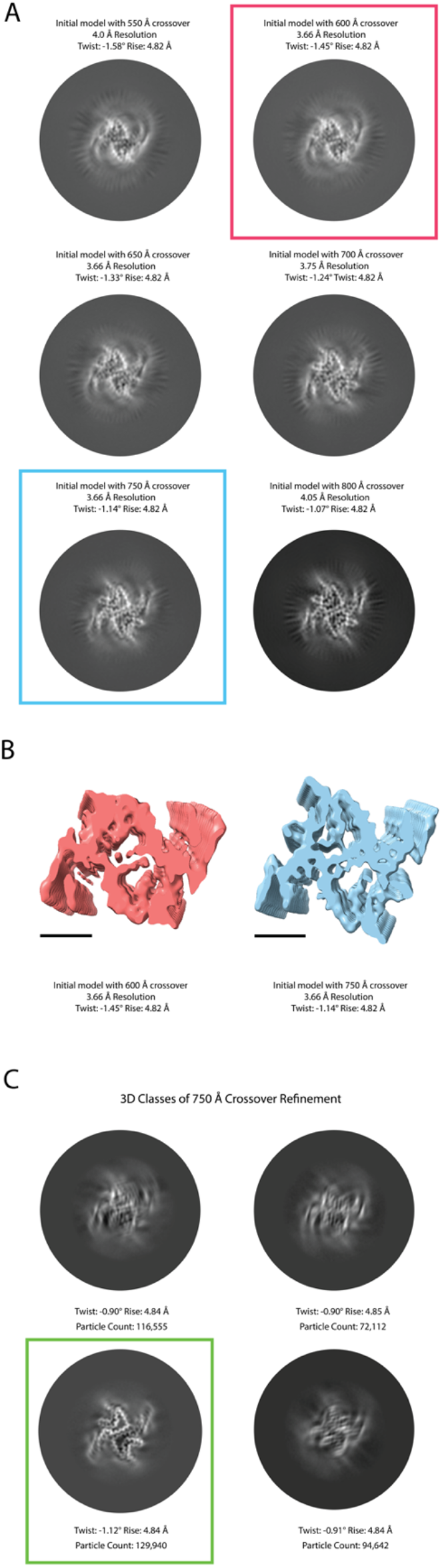
3D Refinement of different crossover distances and 3D classification. A. Cross-sections, resolution, and calculated twist and rise of each initial models after 3D refinement (550-800 Å) (step 23). Red and Blue squares indicate respective electron potential maps for B. B. Cross-section of the electron potential maps refined with 600 Å (Red) and 750 Å (Blue) crossovers. Scale bar, 25 Å. C. 3D Classifications from 750 Å crossover initial model (step 24). Green box indicates selected 3D class used for further refinement (step 25).

**Figure 10.**
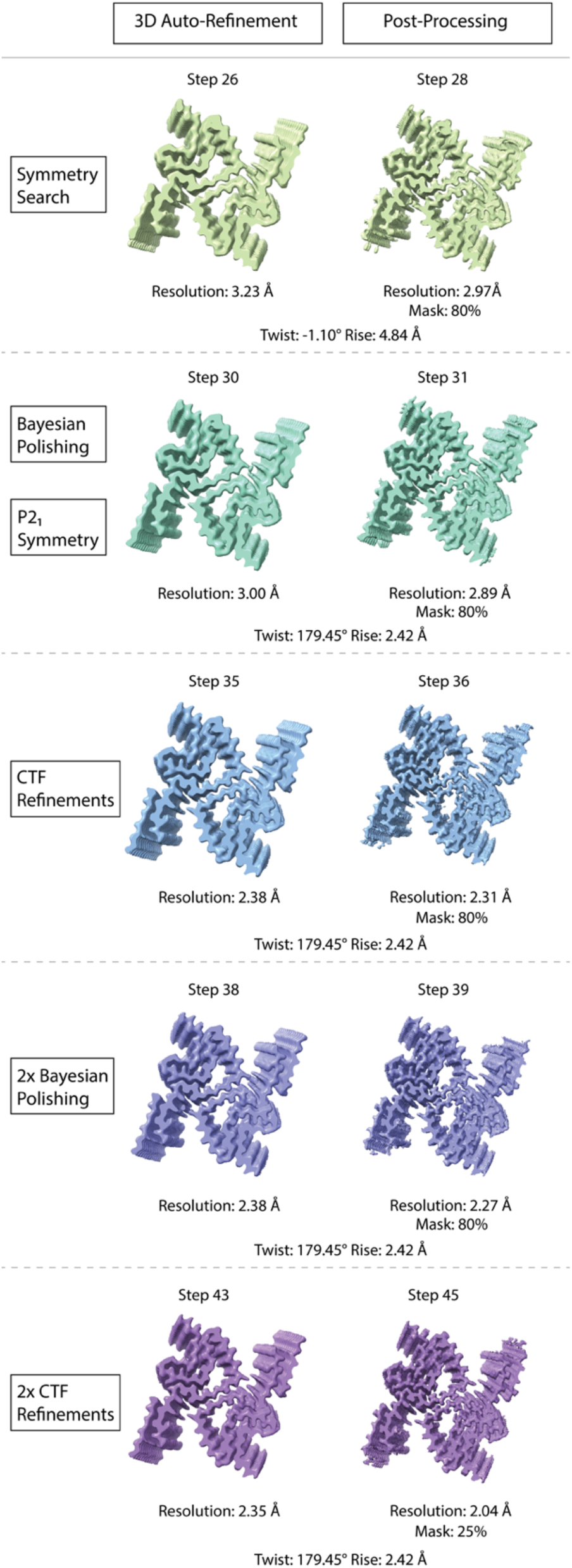
Results of 3D refinements and post-processing steps. The 3D refinements and their corresponding post-processed maps of our processing pipeline are depicted here. The step number, resolution, twist, rise, and mask percentages are displayed for each electron potential map. A description as to whether the electron potential map display is a result of a 3D refinement job or post-processing job is displayed at the top of the figure. The processing workflow incrementally improves maps quality and resolution, resulting in a final map at 2.04 Å resolution.

**Figure 11.**
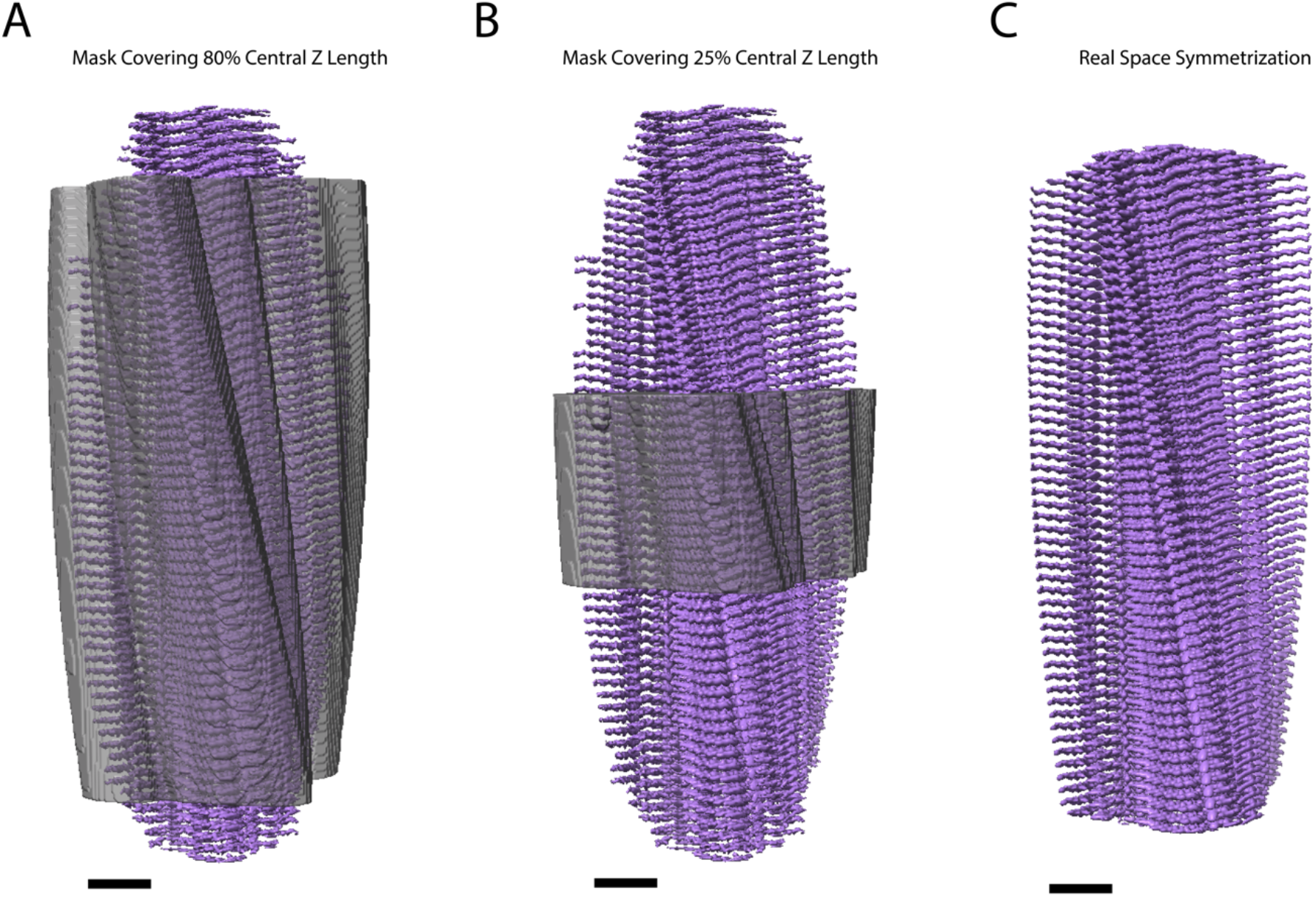
Mask central Z length coverage. A. A mask (gray) covering 80% of the map (purple) along the fibril axis (step 27), used during CTF refinement steps. B. A mask (gray) covering 25% of the map (purple) along the fibril axis (step 44), used in the final post-processing job (step 45). C. Filament after applying real-space symmetrization (step 47) to the edge of the box using the *relion_helix_toolbox* program. Scale bars, 25 Å.

**Figure 12.**
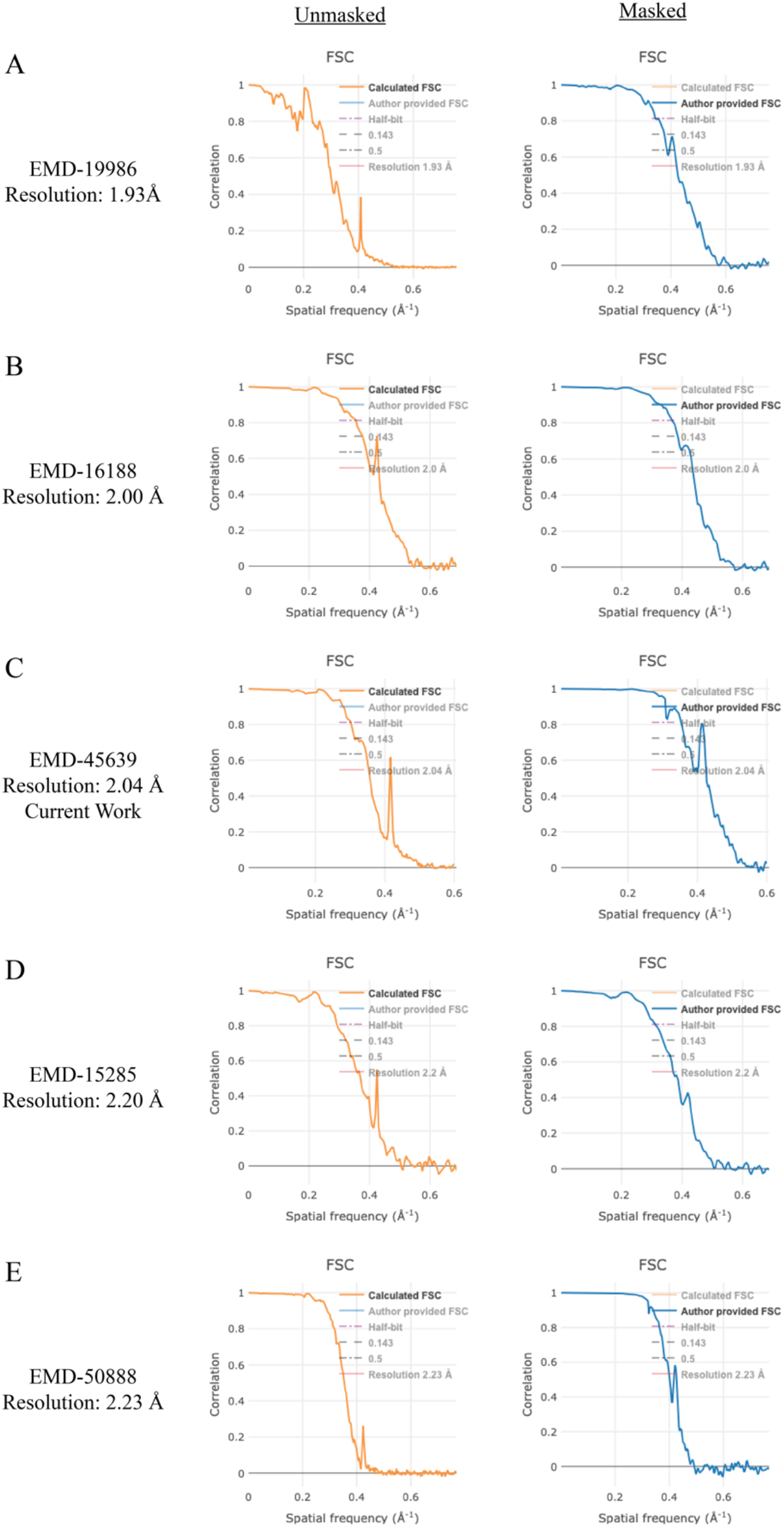
Comparison of Fourier Shell Correlation (FSC) plots of α-synuclein maps deposited to the EMDB resolving to below 2.3 Å. The unmasked FSC plots (calculated FSC from deposited half maps, orange) for the deposited maps display a FSC spike at a spatial frequency of 0.4 Å^-1^ (∼2.4 Å). The masked FSC plots (author provided FSC, blue) dampen this feature.

**Figure 13.**
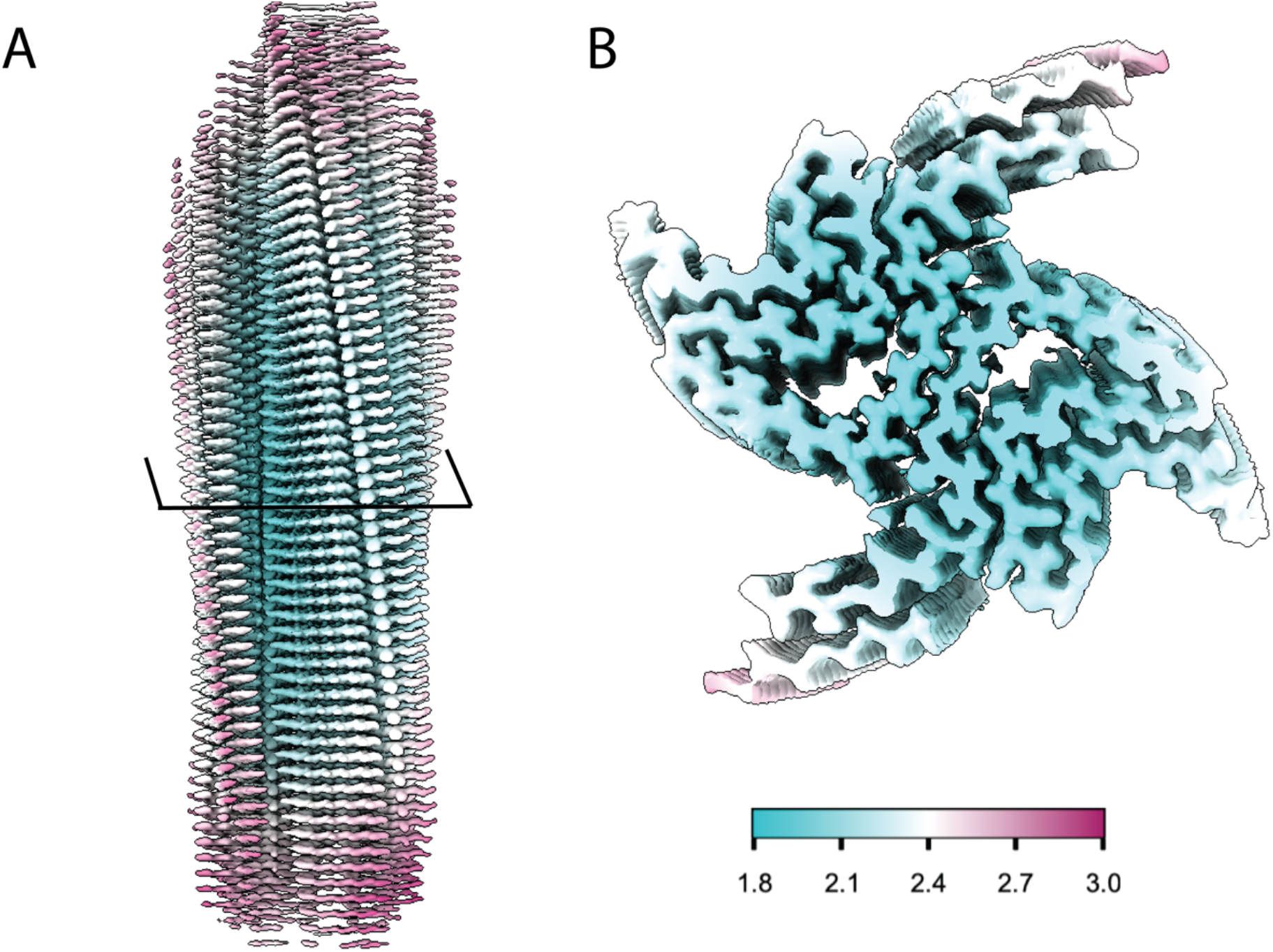
Local resolution map of α-syn fibril from cryo-EM data. A. Local resolution map of filamentous α-syn depicting loss of resolution towards the end of the fibril, with the best resolution located along the central portion of the map. B. Cross-section of the local resolution map of filamentous α-syn showing the best resolved regions of the map are located along the fibril core and protofilament interface. Map resolution key spans from 1.8 Å (cyan) to 3.0 Å (red).

### F. Model building and validation for alpha-synuclein fibrils

There are many methods for building molecular models. Here we used PDB 6H6B as a starting point, the model was fit into the EM map using ChimeraX, then one subunit was rebuilt and refined in Coot. The monomer model was subjected to a round of real-space refinement in Phenix. Then, ChimeraX was used to fit additional refined subunits into the map to generate a multimer model. The multimer model was subjected to final round of real-space refinement in Phenix. We encourage users of this protocol to review tutorials and manuals for ChimeraX, Coot, and Phenix before proceeding with model building [40-42,46,47]. During the modeling process users should use our refined model PDB 9CK3 as a reference. An overview of the entire modeling and validation workflow is provided for reference (Figure 14).

**Figure 14.**
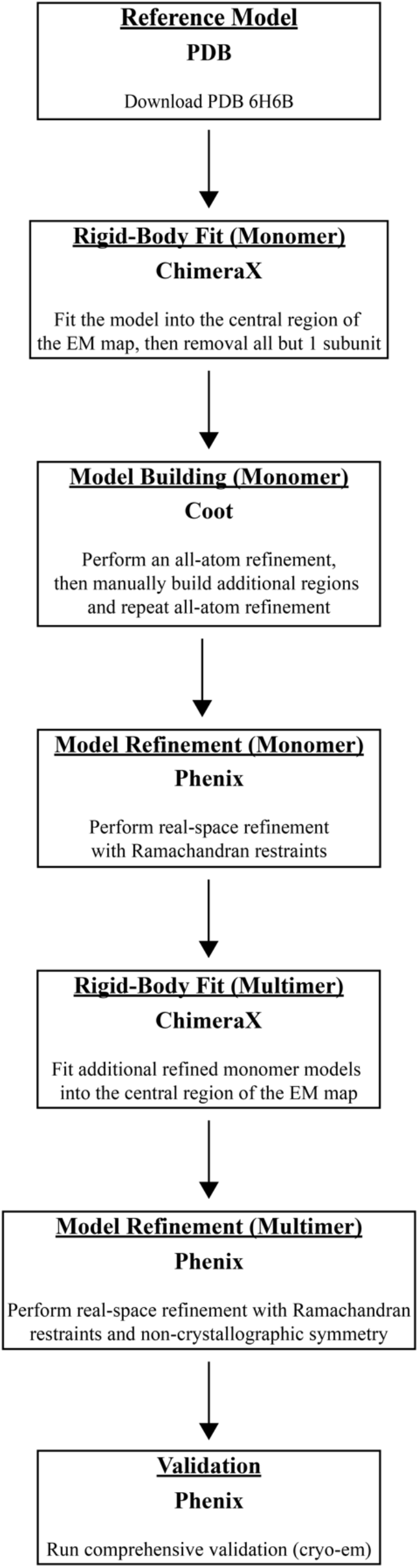
Model building and validation protocol for α-synuclein fibrils. Step by step protocol for building, refining, and validating a α-synuclein fibril molecular model. This protocol uses ChimeraX, Coot, and Phenix in an iterative fashion to improve the molecular model.

1. Download PDB 6H6B by running *open 6h6b* from the ChimeraX command line [12]. This model covers residues 38-95 of the α-synuclein protein and contains 10 monomers displaying the amyloid fold.
2. Open the final *postprocess*.*mrc* file from section G step 45 and under *Volume viewer* set *step size* to *1*.
3. In ChimeraX, use the *Fit* tool to place PDB 6H6B into the central region of the *postprocess*.*mrc* file. You may need to rotate the model to correctly fit the model into the map.
4. Run the command below from the ChimeraX command line to trim the ends of the map for easier visualization of the central region. Then continue fitting until the model is well placed in the map. *view orient; clip front -30 back 30* NOTE: Clipping can be turned off by running *clip off* from the ChimeraX command line.
5. Remove all but 1 monomer from the model by running *split; delete #1*.*2-10* from the ChimeraX command line. We will use Coot to build and refine one subunit and add additional subunits later.
6. Save the file as a PDB, ensure that *Save relative to model* is checked and in the drop down menu select the postprocessed map.
7. In Coot, go to *File → Open Coordinates* and select the PDB file saved in step 6. The go to *File → Open Map* and select the postprocessed map from section G step 45. NOTE: We encourage new users to review the Coot tutorial to become familiar with the software before proceeding (https://www2.mrc-lmb.cam.ac.uk/personal/pemsley/coot/web/tutorial/tutorial.pdf).
8. On the right-hand side is the modeling toolbar, click *Map*, then click *Estimate* to set the map weight, then click *Ok*. NOTE: Click the arrow at the bottom of the modeling toolbar and select *Icons and text* to add the name of each tool to the modeling toolbar.
9. Go to *Refine → All-atom Refine* to improve the fit of the map to the model. When the refinement is done click *Accept* to save the refined atom positions. The refinement will impose geometry restraints and Ramachandran restraints, but it will ignore rotamers. We will handle rotamers in Phenix. NOTE: Use the mouse scroll wheel to adjust the map contour level as necessary throughout this process.
10. The density generated in this protocol allows for modeling of additional residues not resolved in PDB 6H6B, so we need to build additional regions of the model. Go to leucine 38, located at the n-terminus, from the modeling toolbar click *Add Terminal Residue* and click on leucine 38, this will add an alanine residue to the n-terminus. From the modeling toolbar click *Simple Mutate*, then click on alanine 37, a window will appear listing all the amino acids, click *Val (V)* to change alanine 37 to a valine. From the modeling toolbar click *Real Space Refine Zone*, then click on valine 37 and valine 40, this will refine the region between these two residues and improve the fit of the model to the map, then click *Accept* to save the refined atom positions.
11. Repeat step 10, to add glycine 36 to the n-terminus, and lysine 96 and lysine 97 to the c-terminus.
12. Next, build residues 15 to 22 into a well resolved island of density located near the n-terminus. Go to the island of density and rotate the density so the fibril core is oriented towards the top of the screen, this will help minimize the number of movements needed to place the strand into place. Go to *Calculate → Other Modelling Tools → Place Strand Here*, set *Estimated number of residues in strand* to *8* and click *Go*. A strand comprised of 8 alanine residues should now appear, use the *Real Space Refine Zone* tool to improve the fit of the strand into the map.
13. Use the *Simple Mutate* tool to change the alanine strand to the correct residues (V15, V16, A17, A18, A19, E20, K21, T22). Then use the *Real Space Refine Zone* tool to further improve the fit of the strand.NOTE: The n-terminus (i.e. island of density) folds back towards the fibril core adjacent to residues 36 to 44, with residue 15 closest to the fibril core.
14. Click on *Display Manager*, you will see that there are two molecules, one is the PDB that was imported and the second is the new strand that was created. We need to renumber the residues in the new strand, merge the molecules and fix the chain ID. Go to *Edit → Renumber Residues*, under *Renumber Residue Range of Molecule* select the newly generated strand, under *Start Residue* select *N-terminus*, in the *Apply Offset* box provide an integer value to correct for the difference in residue number for the residue that should be valine 15, then click *Apply*. For example, if the valine on the n-terminus of the strand is labeled as V40 then the offset should be -25 to set the valine to residue 15. Click on the n-terminus valine of the strand to verify the numbering is correct.
15. To merge the molecules, Go to *Edit → Merge Molecules*, under *Append/Insert Molecule(s)* select the strand and under *into Molecule* select the original PDB from the drop-down menu, then click *Merge*.
16. Change the chain IDs so both fragments are labeled as chain A. Go to *Edit → Change Chain IDs*, under *Change Chain ID in Molecule* select the file that contains both fragments (i.e. the recently merged molecule), under *From Chain ID* select either chain, under *Using Residue Selection* select *Whole Chain*, under *To Chain ID* set this value to *A*, then click *Apply New Chain ID*. Repeat the process if the second fragment is labeled anything other than chain A. The fragments should now be one molecule labeled as chain A with a dotted line showing the missing residues from residues 23 to 36 that are not resolved.
17. Refine the new molecule that spans residues 15-22 and 37-98. Go to *Refine → All-atom Refine*, if the atoms are well positioned click *Accept*. If not, manually adjust misplaced atoms by dragging the atoms into place and then click *Accept*.
18. Save the coordinates, go to *File → Save Coordinates*, under *Select Molecule Number to Save* select the molecules that was refined in step 17, click *Select Filename* and save the file to the desired location.
19. Open Phenix and setup a new project. NOTE: We encourage new users to review the Phenix tutorial, specifically the real space refinement tutorial, to become familiar with the software before proceeding (https://phenix-online.org/documentation/reference/real_space_refine.html)
20. Under the cryo-Em section select the *Real-space refinement* job. Provide the PDB file from Coot as the model file and the postprocessed file as the map file. Set *Resolution* as determined in the final RELION postprocessing job, in this case the resolution is 2.04 Å. Under the *Refinement Settings* tab, in addition to the default settings ensure *Use secondary structure restraints* and *Ramachandran restraints* is checked, set *Nproc* to 4, click *Rotamers* and under *Fit* select *outliers and poormap*, then click *Run*. Upon completion, the validation report shows that the model statistics are favorable. The *Rotamer outliers (%)* will be slightly elevated due to a salt bridge that forms between lysine 80 and glutamic acid 46, causing lysine 80 to be a rotamer outlier that is supported by the data.
21. In ChimeraX, open the refined model and the postprocessed map. Open the refined model again and now two models are available. Select the second model and use the *Fit* tool to place the second monomer into the opposing protofilament. Repeat the process of opening the refined model and fitting it into a new region of the map. For PDB 9CK3 we built a dodecamer model.
22. Once the desired number of subunits are fitted into the map, run *combine* from the ChimeraX command line to merge the subunits into one model. The command should provide a unique chain ID to each subunit.
23. Repeat step 6 to save the model relative to the postprocessed map.
24. In Phenix, repeat real space refinement as in step 20 with the additional parameter *Ncs constraints* selected. The final validation report shows excellent model statistics with only lysine 80 as a rotamer outlier, as expected. This step can be repeated, if necessary. The model is now ready for structure analysis.

### Additional Validation

This protocol or parts of it has been used and validated in the following research articles:

- Dhavale, et al. [33]. Structure of alpha-synuclein fibrils derived from human Lewy boy dementia tissue. Nature Communications. https://doi.org/10.1038/s41467-024-46832-5.
- Montemayor et al. [48]. Flagellar Structures from the Bacterium Caulobacter crescentus and Implications for Phage φ CbK Predation of Multiflagellin Bacteria. Journal of Bacteriology. https://doi.org/10.1128/jb.00399-20.
- Sanchez et al. [49]. Atomic-level architecture of Caulobacter crescentus flagellar filaments provide evidence for multi-flagellin filament stabilization. bioRxiv. https://doi.org/10.1101/2023.07.10.548443.
- The cryo-EM structure in the manuscript has been validated by our submissions to the PDB (9CK3) and EMDB (EMD-45639). https://doi.org/10.2210/pdb9CK3/pdb.

## Discussion

The fibrilization conditions presented here are specific to one form of *in vitro* assembled α-synuclein fibrils. Extensive optimization of protein purification and fibrilization conditions, testing buffer conditions and incubation parameters, may be necessary to generate different *in vitro* forms. The cryo-EM helical reconstruction methods presented here assume that fibrils are both twisting and are of sufficient length to determine the crossover distance for helical twist estimates. There are cases where fibrils may not twist and thus this workflow would not be amendable to such samples. Finally, structure determination of patient derived fibrils is of high interest, but extraction of fibrils from patient tissue is outside of the scope of the work presented here. Though in theory, the data processing methods presented here should be applicable to these samples.

Cryo-EM data processing is dependent on the sample, data collection instrumentation and parameters used, and computational hardware and software. What we have presented here should provide users with the necessary details for cryo-EM structure determination of a range of amyloid fibrils. We used this approach to generate a cryo-EM map and atomic model of *in vitro* assembled α-synuclein fibrils; and atomic models were deposited in the Protein Data Bank (PDB) under accession 9CK3. Cryo-EM maps, including the final map, half-maps, and mask were deposited in the Electron Microscopy Data Bank (EMDB) under accession EMD-45639.

The work presented here, including sample preparation, NS-TEM, cryo-EM data collection, cryo-EM data processing, and molecular model building serves as a starting point for individuals new to cryo-EM structural analyses of amyloid proteins. For cryo-EM structure determination, new samples will pose their own unique set of challenges, but by first completing the data processing workflow in section E with the EMPIAR dataset under accession EMPIAR-12229, new users will be more adept at troubleshooting new issues.

## Acknowledgements

This work was supported in part by the University of Wisconsin, Madison, the Department of Biochemistry at the University of Wisconsin, Madison, and public health service grants U24 GM139168 to E.R.W, P41GM136463 to C.M.R, and RF1 NS110436 E.R.W. and C.M.R. from the NIH. J.C.S. was supported in part by the Biotechnology Training Program at the University of Wisconsin, Madison, T32 GM135066, the Steenbock Predoctoral Graduate Fellowship administered by the University of Wisconsin-Madison Department of Biochemistry, and the SciMed Graduate Research Scholars Fellowship with support for this fellowship provided by the Graduate School, part of the Office of Vice Chancellor for Research and Graduate Education at the University of Wisconsin-Madison, with funding from the Wisconsin Alumni Research Foundation and the UW-Madison. C.G.B. was supported by the NIH Ruth L. Kirschstein Fellowship, F32 GM149118, from the NIGMS. We are grateful for the critical feedback, guidance, and support provided by Dr. Bryan Sibert, Dr. Matthew Larson, and Ms. Jennifer Scheuren on cryo-EM data collection, data processing, and use of the cryo-EM HPC cluster. We are grateful for the use of facilities and instrumentation at the Cryo-EM Research Center in the Department of Biochemistry at the University of Wisconsin, Madison. We are grateful for the computational resources supplied through the SBGrid Consortium [50].

## Data deposition

The atomic model was deposited in the Protein Data Bank under accession 9CK3. Cryo-EM maps were deposited in the Electron Microscopy Data Bank under accession EMD-45639. The raw micrographs, gain file, and detector MTF file are available on the EMPIAR-12229 database under accession EMPIAR-12229.

## References

1. Baba, M., Nakajo, S., Tu, P. H., Tomita, T., Nakaya, K., Lee, V. M., Trojanowski, J. Q. and Iwatsubo, T. (1998). Aggregation of alpha-synuclein in Lewy bodies of sporadic Parkinson’s disease and dementia with Lewy bodies. Am J Pathol 152(4): 879–884.

2. Spillantini, M. G., Schmidt, M. L., Lee, V. M., Trojanowski, J. Q., Jakes, R. and Goedert, M. (1997). Alpha-synuclein in Lewy bodies. Nature 388(6645): 839–840. 10.1038/42166.

3. Tu, P. H., Galvin, J. E., Baba, M., Giasson, B., Tomita, T., Leight, S., Nakajo, S., Iwatsubo, T., Trojanowski, J. Q. and Lee, V. M. (1998). Glial cytoplasmic inclusions in white matter oligodendrocytes of multiple system atrophy brains contain insoluble alpha-synuclein. Ann Neurol 44(3): 415–422. 10.1002/ana.410440324.

4. Eisenberg, D. S. and Sawaya, M. R. (2017). Structural Studies of Amyloid Proteins at the Molecular Level. Annual Review of Biochemistry 86(Volume 86, 2017): 69–95. 10.1146/annurev-biochem-061516-045104.

5. Burre, J., Sharma, M., Tsetsenis, T., Buchman, V., Etherton, M. R. and Sudhof, T. C. (2010). Alpha-synuclein promotes SNARE-complex assembly in vivo and in vitro. Science 329(5999): 1663–1667. 10.1126/science.1195227.

6. Davidson, W. S., Jonas, A., Clayton, D. F. and George, J. M. (1998). Stabilization of alpha-synuclein secondary structure upon binding to synthetic membranes. J Biol Chem 273(16): 9443–9449. 10.1074/jbc.273.16.9443.

7. Lou, X., Kim, J., Hawk, B. J. and Shin, Y. K. (2017). alpha-Synuclein may cross-bridge v-SNARE and acidic phospholipids to facilitate SNARE-dependent vesicle docking. Biochem J 474(12): 2039–2049. 10.1042/BCJ20170200.

8. Sun, J., Wang, L., Bao, H., Premi, S., Das, U., Chapman, E. R. and Roy, S. (2019). Functional cooperation of alpha-synuclein and VAMP2 in synaptic vesicle recycling. Proc Natl Acad Sci U S A 116(23): 11113–11115. 10.1073/pnas.1903049116.

9. Thapa, K., Khan, H., Kanojia, N., Singh, T. G., Kaur, A. and Kaur, G. (2022). Therapeutic Insights on Ferroptosis in Parkinson’s disease. Eur J Pharmacol 930: 175133. 10.1016/j.ejphar.2022.175133.

10. Serpell, L. C., Berriman, J., Jakes, R., Goedert, M. and Crowther, R. A. (2000). Fiber diffraction of synthetic alpha-synuclein filaments shows amyloid-like cross-beta conformation. Proc Natl Acad Sci U S A 97(9): 4897–4902. 10.1073/pnas.97.9.4897.

11. Spillantini, M. G. and Goedert, M. (2000). The alpha-synucleinopathies: Parkinson’s disease, dementia with Lewy bodies, and multiple system atrophy. Ann N Y Acad Sci 920: 16–27. 10.1111/j.1749-6632.2000.tb06900.x.

12. Guerrero-Ferreira, R., Taylor, N. M., Mona, D., Ringler, P., Lauer, M. E., Riek, R., Britschgi, M. and Stahlberg, H. (2018). Cryo-EM structure of alpha-synuclein fibrils. Elife 7. 10.7554/eLife.36402.

13. Li, Y., Zhao, C., Luo, F., Liu, Z., Gui, X., Luo, Z., Zhang, X., Li, D., Liu, C. and Li, X. (2018). Amyloid fibril structure of alpha-synuclein determined by cryo-electron microscopy. Cell Res 28(9): 897–903. 10.1038/s41422-018-0075-x.

14. Tuttle, M. D., Comellas, G., Nieuwkoop, A. J., Covell, D. J., Berthold, D. A., Kloepper, K. D., Courtney, J. M., Kim, J. K., Barclay, A. M., Kendall, A., et al. (2016). Solid-state NMR structure of a pathogenic fibril of full-length human alpha-synuclein. Nat Struct Mol Biol 23(5): 409–415. 10.1038/nsmb.3194.

15. Lei, Z., Cao, G. and Wei, G. (2019). A30P mutant alpha-synuclein impairs autophagic flux by inactivating JNK signaling to enhance ZKSCAN3 activity in midbrain dopaminergic neurons. Cell Death Dis 10(2): 133. 10.1038/s41419-019-1364-0.

16. Zarranz, J. J., Alegre, J., Gomez-Esteban, J. C., Lezcano, E., Ros, R., Ampuero, I., Vidal, L., Hoenicka, J., Rodriguez, O., Atares, B., et al. (2004). The new mutation, E46K, of alpha-synuclein causes Parkinson and Lewy body dementia. Ann Neurol 55(2): 164–173. 10.1002/ana.10795.

17. Appel-Cresswell, S., Vilarino-Guell, C., Encarnacion, M., Sherman, H., Yu, I., Shah, B., Weir, D., Thompson, C., Szu-Tu, C., Trinh, J., et al. (2013). Alpha-synuclein p.H50Q, a novel pathogenic mutation for Parkinson’s disease. Mov Disord 28(6): 811–813. 10.1002/mds.25421.

18. Lesage, S., Anheim, M., Letournel, F., Bousset, L., Honore, A., Rozas, N., Pieri, L., Madiona, K., Durr, A., Melki, R., et al. (2013). G51D alpha-synuclein mutation causes a novel parkinsonian-pyramidal syndrome. Ann Neurol 73(4): 459–471. 10.1002/ana.23894.

19. Pasanen, P., Myllykangas, L., Siitonen, M., Raunio, A., Kaakkola, S., Lyytinen, J., Tienari, P. J., Poyhonen, M. and Paetau, A. (2014). Novel alpha-synuclein mutation A53E associated with atypical multiple system atrophy and Parkinson’s disease-type pathology. Neurobiol Aging 35(9): 2180 e2181–2185. 10.1016/j.neurobiolaging.2014.03.024.

20. Polymeropoulos, M. H., Lavedan, C., Leroy, E., Ide, S. E., Dehejia, A., Dutra, A., Pike, B., Root, H., Rubenstein, J., Boyer, R., et al. (1997). Mutation in the alpha-synuclein gene identified in families with Parkinson’s disease. Science 276(5321): 2045–2047. 10.1126/science.276.5321.2045.

21. Yoshino, H., Hirano, M., Stoessl, A. J., Imamichi, Y., Ikeda, A., Li, Y., Funayama, M., Yamada, I., Nakamura, Y., Sossi, V., et al. (2017). Homozygous alpha-synuclein p.A53V in familial Parkinson’s disease. Neurobiol Aging 57: 248 e247–248 e212. 10.1016/j.neurobiolaging.2017.05.022.

22. Frey, L., Ghosh, D., Qureshi, B. M., Rhyner, D., Guerrero-Ferreira, R., Pokharna, A., Kwiatkowski, W., Serdiuk, T., Picotti, P., Riek, R., et al. (2023). On the pH-dependence of α-synuclein amyloid polymorphism and the role of secondary nucleation in seed-based amyloid propagation.

23. Mehra, S., Gadhe, L., Bera, R., Sawner, A. S. and Maji, S. K. (2021). Structural and Functional Insights into α-Synuclein Fibril Polymorphism. Biomolecules 11(10): 1419.

24. Peelaerts, W., Bousset, L., Van der Perren, A., Moskalyuk, A., Pulizzi, R., Giugliano, M., Van den Haute, C., Melki, R. and Baekelandt, V. (2015). α-Synuclein strains cause distinct synucleinopathies after local and systemic administration. Nature 522(7556): 340–344. 10.1038/nature14547.

25. Marotta, N. P., Ara, J., Uemura, N., Lougee, M. G., Meymand, E. S., Zhang, B., Petersson, E. J., Trojanowski, J. Q. and Lee, V. M. Y. (2021). Alpha-synuclein from patient Lewy bodies exhibits distinct pathological activity that can be propagated in vitro. Acta Neuropathologica Communications 9(1): 188. 10.1186/s40478-021-01288-2.

26. Uemura, N., Marotta, N. P., Ara, J., Meymand, E. S., Zhang, B., Kameda, H., Koike, M., Luk, K. C., Trojanowski, J. Q. and Lee, V. M. Y. (2023). α-Synuclein aggregates amplified from patient-derived Lewy bodies recapitulate Lewy body diseases in mice. Nature Communications 14(1): 6892. 10.1038/s41467-023-42705-5.

27. Yang, Y., Shi, Y., Schweighauser, M., Zhang, X., Kotecha, A., Murzin, A. G., Garringer, H. J., Cullinane, P. W., Saito, Y., Foroud, T., et al. (2022). Structures of α-synuclein filaments from human brains with Lewy pathology. Nature 610(7933): 791–795. 10.1038/s41586-022-05319-3.

28. Riek, R. and Eisenberg, D. S. (2016). The activities of amyloids from a structural perspective. Nature 539(7628): 227–235. 10.1038/nature20416.

29. Jarrett, J. T. and Lansbury, P. T., Jr. (1992). Amyloid fibril formation requires a chemically discriminating nucleation event: studies of an amyloidogenic sequence from the bacterial protein OsmB. Biochemistry 31(49): 12345–12352. 10.1021/bi00164a008.

30. Tornquist, M., Michaels, T. C. T., Sanagavarapu, K., Yang, X., Meisl, G., Cohen, S. I. A., Knowles, T. P. J. and Linse, S. (2018). Secondary nucleation in amyloid formation. Chem Commun (Camb) 54(63): 8667–8684. 10.1039/c8cc02204f.

31. Tanaka, G., Yamanaka, T., Furukawa, Y., Kajimura, N., Mitsuoka, K. and Nukina, N. (2019). Sequence- and seed-structure-dependent polymorphic fibrils of alpha-synuclein. Biochimica et Biophysica Acta (BBA) - Molecular Basis of Disease 1865(6): 1410–1420. 10.1016/j.bbadis.2019.02.013.

32. Strohäker, T., Jung, B. C., Liou, S.-H., Fernandez, C. O., Riedel, D., Becker, S., Halliday, G. M., Bennati, M., Kim, W. S., Lee, S.-J., et al. (2019). Structural heterogeneity of α-synuclein fibrils amplified from patient brain extracts. Nature Communications 10(1): 5535. 10.1038/s41467-019-13564-w.

33. Dhavale, D. D., Barclay, A. M., Borcik, C. G., Basore, K., Berthold, D. A., Gordon, I. R., Liu, J., Milchberg, M. H., O’Shea, J. Y., Rau, M. J., et al. (2024). Structure of alpha-synuclein fibrils derived from human Lewy body dementia tissue. Nature Communications 15(1): 2750. 10.1038/s41467-024-46832-5.

34. Naiki, H., Higuchi, K., Hosokawa, M. and Takeda, T. (1989). Fluorometric determination of amyloid fibrils in vitro using the fluorescent dye, thioflavine T. Analytical Biochemistry 177(2): 244–249. 10.1016/0003-2697(89)90046-8.

35. Kimanius, D., Dong, L., Sharov, G., Nakane, T. and Scheres, S. H. W. (2021). New tools for automated cryo-EM single-particle analysis in RELION-4.0. Biochem J 478(24): 4169–4185. 10.1042/bcj20210708.

36. Zheng, S. Q., Palovcak, E., Armache, J.-P., Verba, K. A., Cheng, Y. and Agard, D. A. (2017). MotionCor2: anisotropic correction of beam-induced motion for improved cryo-electron microscopy. Nat Methods 14(4): 331–332. 10.1038/nmeth.4193.

37. Zhang, K. (2016). Gctf: Real-time CTF determination and correction. J Struct Biol 193(1): 1–12. 10.1016/j.jsb.2015.11.003.

38. Bepler, T., Morin, A., Rapp, M., Brasch, J., Shapiro, L., Noble, A. J. and Berger, B. (2019). Positive-unlabeled convolutional neural networks for particle picking in cryo-electron micrographs. Nat Methods 16(11): 1153–1160. 10.1038/s41592-019-0575-8.

39. Scheres, S. H. W. (2020). Amyloid structure determination in RELION-3.1. Acta Crystallogr D Struct Biol 76(Pt 2): 94–101. 10.1107/S2059798319016577.

40. Pettersen, E. F., Goddard, T. D., Huang, C. C., Meng, E. C., Couch, G. S., Croll, T. I., Morris, J. H. and Ferrin, T. E. (2021). UCSF ChimeraX: Structure visualization for researchers, educators, and developers. Protein Sci 30(1): 70–82. 10.1002/pro.3943.

41. Goddard, T. D., Huang, C. C., Meng, E. C., Pettersen, E. F., Couch, G. S., Morris, J. H. and Ferrin, T. E. (2018). UCSF ChimeraX: Meeting modern challenges in visualization and analysis. Protein Sci 27(1): 14–25. 10.1002/pro.3235.

42. Meng, E. C., Goddard, T. D., Pettersen, E. F., Couch, G. S., Pearson, Z. J., Morris, J. H. and Ferrin, T. E. (2023). UCSF ChimeraX: Tools for structure building and analysis. Protein Sci 32(11): e4792. 10.1002/pro.4792.

43. Fitzpatrick, A. W. P., Falcon, B., He, S., Murzin, A. G., Murshudov, G., Garringer, H. J., Crowther, R. A., Ghetti, B., Goedert, M. and Scheres, S. H. W. (2017). Cryo-EM structures of tau filaments from Alzheimer’s disease. Nature 547(7662): 185–190. 10.1038/nature23002.

44. Yang, Y., Garringer, H. J., Shi, Y., Lovestam, S., Peak-Chew, S., Zhang, X., Kotecha, A., Bacioglu, M., Koto, A., Takao, M., et al. (2023). New SNCA mutation and structures of alpha-synuclein filaments from juvenile-onset synucleinopathy. Acta Neuropathol 145(5): 561–572. 10.1007/s00401-023-02550-8.

45. Egelman, E. H. (2024). Helical reconstruction, again. Current Opinion in Structural Biology 85: 102788. 10.1016/j.sbi.2024.102788.

46. Emsley, P., Lohkamp, B., Scott, W. G. and Cowtan, K. (2010). Features and development of Coot. Acta Crystallogr D 66: 486–501. 10.1107/S0907444910007493.

47. Liebschner, D., Afonine, P. V., Baker, M. L., Bunkoczi, G., Chen, V. B., Croll, T. I., Hintze, B., Hung, L. W., Jain, S., McCoy, A. J., et al. (2019). Macromolecular structure determination using X-rays, neutrons and electrons: recent developments in Phenix. Acta Crystallogr D Struct Biol 75(Pt 10): 861–877. 10.1107/S2059798319011471.

48. Montemayor, E. J., Ploscariu, N. T., Sanchez, J. C., Parrell, D., Dillard, R. S., Shebelut, C. W., Ke, Z., Guerrero-Ferreira, R. C. and Wright, E. R. (2021). Flagellar Structures from the Bacterium Caulobacter crescentus and Implications for Phage varphi CbK Predation of Multiflagellin Bacteria. J Bacteriol 203(5). 10.1128/JB.00399-20.

49. Sanchez, J. C., Montemayor, E. J., Ploscariu, N. T., Parrell, D., Baumgardt, J. K., Yang, J. E., Sibert, B., Cai, K. and Wright, E. R. (2023). Atomic-level architecture of Caulobacter crescentus flagellar filaments provide evidence for multi-flagellin filament stabilization. bioRxiv. 10.1101/2023.07.10.548443.

50. Morin, A., Eisenbraun, B., Key, J., Sanschagrin, P. C., Timony, M. A., Ottaviano, M. and Sliz, P. (2013). Collaboration gets the most out of software. Elife 2: e01456. 10.7554/eLife.01456.

